# Effect of structure in ionized albumin – based nanoparticle: Characterisation, Emodin interaction, and *in vitro* cytotoxicity

**DOI:** 10.1101/631788

**Authors:** Macarena Siri, Maria Julieta Fernandez Ruocco, Estefanía Achilli, Malvina Pizzuto, Juan F. Delgado, Jean-Marie Ruysschaert, Mariano Grasselli, Silvia del V. Alonso

**Affiliations:** Laboratorio de Biomembranas (LBM), Departamento de Ciencia y Tecnología, Universidad Nacional de Quilmes, IMBICE-CONICET-CICPBA; Laboratorio de Materiales Biotecnológicos (LaMaBio), Departamento de Ciencia y Tecnología, Universidad Nacional de Quilmes, IMBICE-CONICET-CICPBA; Instituto de Biofísica Carlos Chagas Filho, Universidade Federal do Rio de Janeiro, Brazil; Laboratorio de Obtención, Modificación, Caracterización y Evaluación de Materiales (LOMCEM), Conicet, Universidad Nacional de Quilmes; Laboratory for the Structure and Function of Biological Membranes, Center for Structural Biology and Bioinformatics, Université Libre de Bruxelles, CP 206/02, Bd du Triomphe, 1050 Brussels, Belgium

**Keywords:** Bovine Serum Albumin (BSA), BSA-nanoparticle (BSA NP), Emodin, drug delivery

## Abstract

A γ–irradiated bovine albumin serum based nanoparticle was characterised structurally, and functionally. The nanoparticle was characterised by A.F.M, D.L.S, zeta potential, T.E.M., gel-electrophoresis, spectroscopy (UV-Vis, Fluorescence, FT-IR, and CD). Its stability was studied under adverse experimental conditions: pH values, chaotropic agents, and ionic strength and stability studies against time were mainly carried out by fluorescence spectroscopy following the changes in the tryptophan environment in the nanoparticle. Its function was studied by the interaction of the NP with the hydrophobic drug Emodin was studied. The binding and kinetic properties of the obtained complex were tested by biophysical methods as well as its toxicity in tumour cells.

According to its biophysics, the nanoparticle is a spherical nanosized vehicle with a hydrodynamic diameter of 70 nm. Data obtained describe the nanoparticle alone as nontoxic for cancer cell lines. When combined with Emodin, the bioconjugate proved to be more active on MCF-7 and PC-3 cancer cell lines than the nanoparticle alone. No haemolytic activity was found when tested against *ex vivo* red blood cells. The stability of the albumin nanoparticle is based on a competition between short-range attraction forces and long-range repulsion forces. The nanoparticle showed similar behaviour as albumin against pH while improving its stability in urea and tween 80. It was stable up to 15 days and presented no protein degradation in solutions up to 2 M salt concentration. Significantly, the albumin aggregate preserves the main activity-function of albumin and improved characteristics as an excellent carrier of molecules.

**Graphical Abstract:** 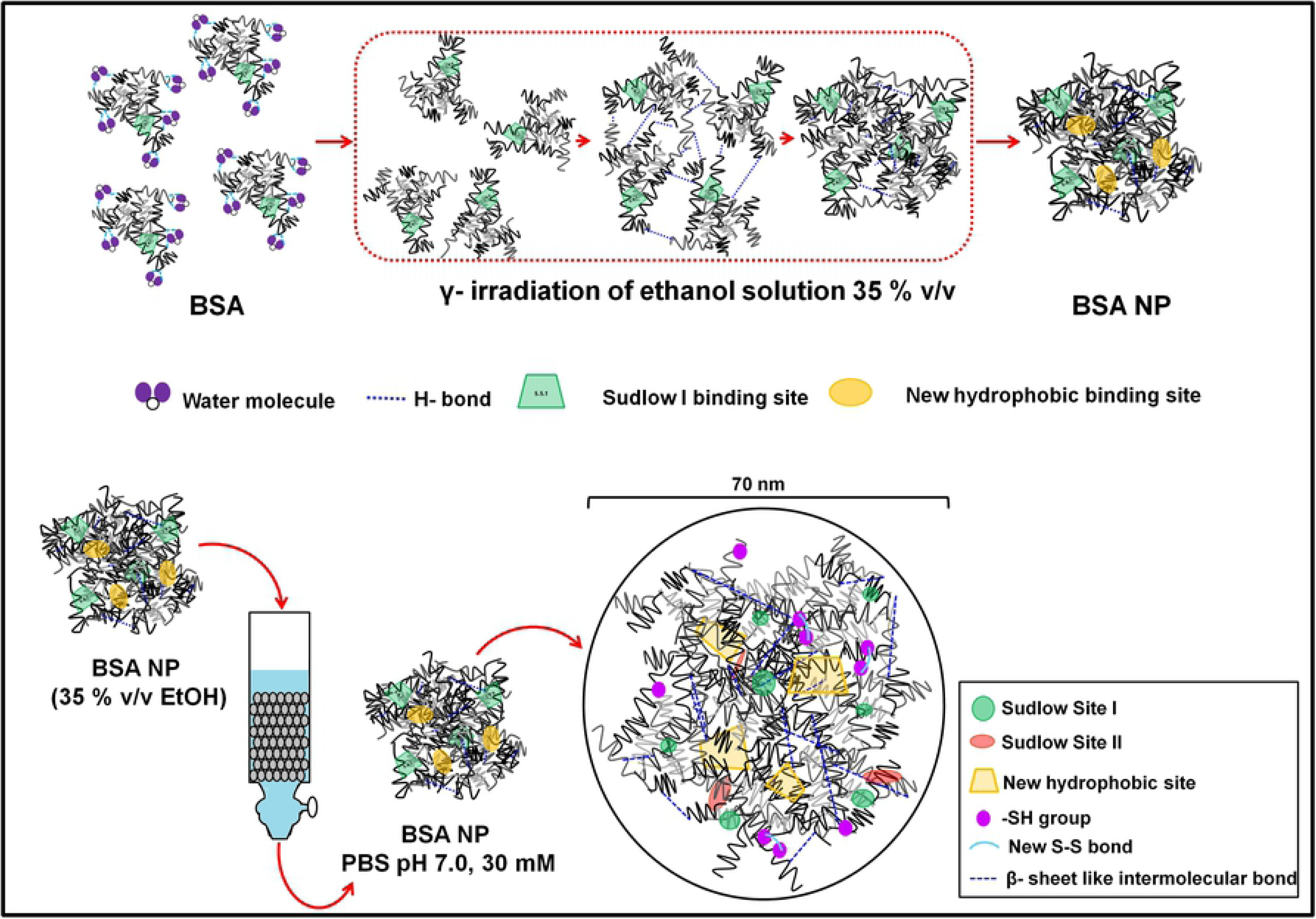

## Introduction

Protein-based nanoparticles constitute an alternative to classical protein-ligand complexes as drug delivery systems. They also display interesting biophysical and functional properties based on multi-site localized specific binding sites of the protein surface [1].

When designing and preparing a protein nanoparticle (NP), the preservation of its primary function is highly essential. The challenge is not to let the preparation method influence the primary function of the protein, despite the nanoparticle undergoing small changes in its secondary structure. Proteins forming part of the NP can face substantial challenges to their activity due to structural alteration during the NP preparation process. If the alteration results in loss of functionality, then following protein modification might be required. However, if the NP is designed in the right way, it could significantly enhance the delivery of the desired substance [2]. Current techniques based mainly on chemical methods are employed for the synthesis of protein nanoparticles [3]. A highlighted feature of proteins is their capacity to aggregate in a size-dependent manner according to the ethanol concentration used. The aggregation eases the tailoring into desirable functional size-stable NPs for *in vivo* penetration. The preparation methods in ethanol solution and lyophilisation of the BSA NP were published in previous works [3][4][5]. In this structural study, we give a complete biophysical characterisation of the gamma-irradiated albumin-based nanoparticle, where DLS size and microscopy was covered before [4]. In order to propose it as a suitable drug delivery system, not only structural but also functional studies is given. As BSA is known as an excellent carrier for drugs, it is expected that the NP should be able to bind to antitumor drugs. In this work, we succeeded in preparing an NP by using γ irradiation as a crosslinker in order to avoid residual toxic organic secondary products and help control the size protein NP [3][6]. NP destined to be delivery vehicles have to be designed with particular care in its size, charge and surface chemistry as these characteristics are of great importance as they strongly affect their behaviour once in blood circulation [7]. The design of a new nanoparticle should:

1. Present stability in a biological suspension (pH around 7.4) and bear high salt concentration in a solution;
2. Have the ability to avoid an immediate uptake by the reticuloendothelial system;
3. Be able to provide functional surface groups for further decoration, and
4. Be able to resist stress like low pH values if administrated orally [8]
5. Be able to maintain stable conformation during storage [9]
6. Be able to avoid aggregation

Therefore, the behaviour under different conditions comes out as an essential study item in the field of nanoparticle study.

One of the most popular proteins used for NP preparation is albumin. The BSA tertiary structure is composed of three domains: I, II and III. Each domain is formed by six helices and two subdomains which are named IAB, IC, IIAB, IIC, IIIAB, and IIIC. Both domains II and III are the ones that present the primary binding sites of the protein. It also contains aromatic residues shared by two or more of these sites [9]. Both binding sites, named Sudlow sites share a tryptophan (Trp – 212). This aminoacid limits the accessibility of the solvent in IIA binding site and takes part in the hydrophobic packing interactions at the IIA- IIIA interface [10]. Therefore, this Trp is used to monitor the conformational changes in the protein by spectroscopic techniques. Sudlow sites 1 and two display affinity with a wider variety of elements. Of the two given sites, site 1 has the highest affinity with hydrophobic drugs; when bound, a displacement of the Trp 212 occurs [9] [11]. It presents appropriate binding sites for hydrophobic drugs, given its biodegradability to natural products, non-toxic and immunogenicity the reason for multiple drugs transports [9][12][13]. Moreover, albumin belongs to a multigenetic family which includes α-fetoprotein and vitamin D-binding protein. It is the most soluble protein in vertebrates, as well as, the most abundant and non-toxic carrier. It has in its structure 18-21 Tyr, depending on the type of albumin. Given the presence of Trp 212 and 143 in BSA, this makes it suitable for spectrophotometric studies [11].

In this structural study, we give a complete biophysical characterisation of the gamma-irradiated albumin-based nanoparticle, where DLS size and microscopy was covered before [4]. In order to propose it as a suitable drug delivery system, not only structural but also functional studies are given. As BSA is known as an excellent carrier for drugs, it is expected that the NP should be able to bind to antitumour drugs. This study aim is to characterise a bovine serum albumin nanoparticle (BSA NP). This nanoparticle presents the advantage of only having BSA in its composition. Therefore, if its stable, the potentiality of this new technology would rise as toxicity levels are expected to be low, and its preparation, easy and reproducible [3][4][13]. The stability experiments at different pHs and chaotropic were carried out mainly by fluorescence spectroscopy.

In some cases, the findings of the NP behaviour were contrasted with that of the albumins. We also aim at the biophysical characterisation of the NP bioconjugate Emodin (E) / BSA NP (BSA NPE) as a drug delivery system. As a result, a confirmation of a stable, non-toxic NP with small structural changes was obtained. The toxicity effect *in vitro* in cell lines like MCF7 and PC3 and in an *in vivo* model (zebrafish) was evaluated on the free drug, free NP and the bioconjugate BSA NPE. The results presented here will help understand the nanoparticles–protein-based drug interactions and the role that BSA nanoparticles may play in future biomedical and pharmacological applications.

The drug used in this work is emodin (3-methyl-1,6,8-trihydroxyanthraquinone). The drug is an orange natural anthraquinone that has been commonly used for its anti-inflammatory and laxative effect. Emodin (E) is a molecule with a structure similar to the anthracene, which has shown in recent years an antitumor effect. It features as an antitumor drug, producing apoptosis by the caspases way in cell lines derived from tumours such as HL60 [2] and HeLa [14][15][16]. It presents an acid-alkaline equilibrium in aqueous solutions, varying back and forward from neutral species, monoanionic and dianionic species [9][11]. As it is this drug has been studied in recent years because of its many properties, and attempts have been made to diminish its secondary effects by different drug delivery systems [4][17][18]. The most relevant feature of E is its ability to bind to proteins forming complexes. Hence, the reason for choosing said drug in this study [2][11].

## 2. Experimental Section

### 2.1. Materials

BSA was purchased from Sigma-Aldrich (A7906-100G Lot SLBC371V), 98% agarose gel electrophoresis, lyophilized powder (Buenos Aires, Argentina). Ethanol was purchased from Biopack (Zarate, Buenos Aires, Argentina) and acetone was obtained from Anedra (Research AG, Tigre, Buenos Aires, Argentina). Sephadex G-200 column for chromatography was obtained from Pharmacia Fine Chemicals (New Jersey, United States of America). Centrifugation tubes were Amicon Ultra, Centrifugal Filtres – 0.5 ml 3K, Ultracel-30 kDa Membrane from Merck Millipore Corporation (Darmstadt, Germany). Urea was purchased from MERCK (Darmtastd, Alemania), SDS was purchased from Anedra (Tigre, Buenos Aires, Argentina) and Tween 80% from FLUKA (Sigma-Aldrich-Buenos Aires, Argentina). All corresponding buffers were PBS analytical grade made.

### 2.2. Nanoparticle preparation

BSA nanoparticles (BSA NP) were obtained by gamma irradiation according to Soto Espinoza et al., 2012 [3]. The solution in which the BSA NP was obtained was a buffer/ ethanol solution. The buffer corresponded to 30 mM phosphate buffer solution (PBS) at a pH 7.0, and the ethanol concentration was 35 % v/v.

Briefly, BSA molecules in a solution of buffer/ ethanol solution were gamma irradiated by a ^60^Co source (at PISI CNEA-Ezeiza, Argentina), at a dose rate of 1 kGy/h (lowest dose set at two kGy and upper limit dose set at 10 kGy). The sample temperature during irradiation was between 5-10 °C. After, samples were eluted in a molecular exclusion column to exchange the ethanol solution for 30 mM PBS pH 7.

### 2.3 Nanoparticle characterisation

All experiments were referred to BSA (MW 66 kDa) of an initial concentration of 4.5 to 450 µM, which corresponds to concentrations of nanoparticle between 5.5 nM to 555 nM. The following concentrations were calculated based on the BSA NP molecular weight (MW 148 MDa) obtained from TEM (spherical shaped NPs) and DLS nanoparticle size of 70 nm. The DLS and Z Potential measures were taken with 90 Plus/Bi-MAS particle size analyzer as in Siri et al. 2016 [5].

#### 2.3.1. Size molecular exclusion

BSA NP was purified by size exclusion chromatography using a Sephadex G-200 column. Eluted fractions were followed by UV-Vis (λ= 280 nm). The hydrodynamic size of the eluted peak was determined through Dynamic Light Scattering analyses.

#### 2.3.2. SDS – PAGE

Different BSA NP obtained from different processes were tested in an SDS-PAGE. A total of 30 μl of each BSA NP were diluted in distilled water and 10 μl of sample buffer reaching a final concentration of 5 μM of protein. Samples were heated up for 5 minutes. Aliquots of 18 μl of sample and protein markers were loaded on an 8 % polyacrylamide gel and 5 % stacking gel. A Biopack Electrophoresis Unit was used to run the SDS – PAGE at a constant voltage of 8 mA. After, the gel was stained using Coomassie blue. For gel analysis, ImageJ Gel Analyses was used.

#### 2.3.3. Transmission electron microscopy (T.E.M)

A Philips high-resolution transmission electron microscopy EM 10A/B 60 KV was used for sample morphology observation. The BSA NP buffer solution was diluted 1/100 for this study (final concentration of 5 nM). Samples were stained with uranyl acetate before the microscopies. A total of 20 microscopies was taken over different fields of the sample. The NP selection was carried out by identification of them based on round shape and definition of the software for analysis used was ImageJ (FiJI). A counting tool from the software was used in order to count the NP in the microscopies and then the profile plot tool was used to describe the size of each NP found.

#### 2.3.4. Atomic force microscopy (A.F.M.)

The BSA NP sample was purified by exclusion chromatography in PBS. A total of 20 µl of the samples were deposited on freshly cleaved mica. After, it was dried for 10 min with nitrogen flux. Images were obtained using a Dimension Icon in Peak Force QNM (PFQNM) (Bruker®). Measures were made with rectangular silicon tip with a nominal spring constant 42 N/m and tip radius of 12 nm. This microscopy took place in the Institute of Biophysics, Universidade Federal do Rio de Janeiro, Rio de Janeiro, Brazil.

#### 2.3.5. UV-Vis spectrophotometry

The absorbance profile of the BSA NP was measured in a Nano-Drop 1000 Thermo Scientific spectrophotometer, with a path length of 0.1 cm. Samples were diluted to a concentration of 5.5 nM in 30 mM PBS (pH 7. 0). A total of 4 µl of each sample was needed for the measure. The profiles obtained went from 240 – 400 nm. Plotting of the measurements and data analysis were done using GraphPad Prism v.5.

#### 2.3.6. Fluorescence spectrophotometry

The fluorescence emission profile of the BSA NP was measured in an S2 Scinco Fluorspectrophotometer. The excitation and emission slits were open at a 1.5 nm distance. BSA NP samples (18 nM) were excited at a wavelength of 295 nm, where emission range was 300 - 400 nm. An ml per sample in 30 mM PBS (pH 7.0) was used in the measurements. Plotting of the measurements and data analysis were done using GraphPad Prism v.5.

#### 2.3.7. Circular Dichroism (C.D.)

The CD profile of the BSA NP was measured in a Jasco 810 spectropolarimeter equipped with a Peltier cell device for temperature control (Jasco Corporation, Japan), at room temperature with near and far dichroic signal range (240 – 400 nm and 180 – 280 nm, respectively). A quartz cell of 500 μl was needed for the experiment. Concentrations varied according to the signal range desired to record: for the near – CD BSA NP (555 nM), for the far-CD BSA NP (37 nM). Resolution of the samples was of 0.1 nm. Plotting of the measurements and data analysis were done using GraphPad Prism v.5.

#### 2.3.8. FT-IR Spectroscopy

The infrared profile of the BSA NP was measured in an FT-IR Affinity1-Shimadzu ATR with a laser lamp with an absorbance HAPP-GENZEL of 324. Measurements ran wavelength from 400 to 4000 cm^−1^, with 80 scans per sample and a resolution of 1 cm^−1^. The cell needed, was of SeZn with 45° of inclination. Samples were blown-dried until a film was formed to obtain the BSA NP spectrum free of IR-water signals with The concentration of the sample was 55 nM. The blank was subtracted by software. The spectra were processed and analysed with the IR solution software (v. 1.50), provided by the manufacturer, and by Prism GraphPad v.5.

#### 2.3.9. Free thiols groups’ detection

Samples of BSA NP were diluted 1/10 to a concentration of 0.55 µM for the experiment. The experimental conditions were carried out as described in Grasse et al., 1967 [6] and modified as in Achilli et al., 2015 [4]. Briefly, the absorbance of the sample at 280 nm were measured. A solution of Ellman’s reagent (DTNB) 0.1 M PBS at a 3.9 mg/ml concentration was added to each sample. The incubation lasted for 15 minutes, and then the absorbance at 425 mn was measured again. In order to know how many free thiols we had per NP, we followed the equation,

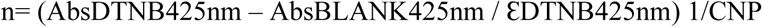

where n is the number of moles of thiols per BSA NP, and ƐDTNB425nm is 12400 (mol cm^−1^). CNP represents the concentration of BSA NP in the sample.

#### 2.3.10. Free amino group’s detection

Solutions of BSA NP (18 nM) were used for these measurements. Experimental protocol followed Habeeb et al., 1966 [19] method for free amino determination. Briefly, to an ml of sample was added 1 ml of 4% NaHCO3, pH 8.5 and 1 ml of 0.1 % TNBS. Then, was incubated for two hours at 42 °C, in the dark. After, 1 ml of 10 % SDS was added to make the NP soluble together with 500 µl of 1N HCl to prevent its precipitation. Samples were measured before and after treatment for its absorbance at 335 nm.

The ratio between TNBS and BSA NP was estimated as [TNBS]/ [BSA NP]. In order to know the TNBS concentration in the sample, a calibration curve was done.

#### 2.3.11. Determination of surface carbonyl groups (protein oxidation)

Solutions of BSA NP (55 nM) were used in this experiment. The procedure followed Brady’s reaction protocol [20]. Briefly, 800 µl of 10 mM DNP in 2 M HCl was added to 200 µl of BSA NP. The incubation lasted an hour at room temperature and in the dark. Every 15 minutes samples were vortexed to promote great mixing between the solutions. After incubation finished, an ml of 20 % w/v of TCA was added, and samples were then incubated for 5 minutes in ice. Once incubation finished, the samples were centrifugated at 10000 g at 4 °C. The pellet from the centrifugation was resuspended in 500 µl of 6 M guanidine. Another centrifugation at 4 °C for 10 minutes (10000 g) took place before measuring the samples by UV-vis absorbance at 370 nm. For the calculation on the ratio carbonyl groups/ NP, the same steps from 2.4.10 were performed.

#### 2.3.12. TGA

BSA NP and BSA samples in miliQ aqueous solution were normalised to an absorbance of 0.371 for this experiment. Samples were lyophilised overnight in order to eliminate any trace of water for the experiment as described in Siri et al., 2016. Approximately 4 mg of each sample was placed in sample pan and submitted to a heating ramp from 30 °C to 920 °C at 10 °C min-1 in a thermogravimetric balance (TA Q500, TA Instruments, United States). The gas used in the sample was air at 40 mL min-1 and nitrogen at 60 mL min-1 as balance gas. Weight percentage was registered in function of temperature. From original data, derivative of weight concerning temperature and temperature at 5% weight loss (T95) were calculated. Three independent samples of BSA NP and BSA were analysed.

### 2.4 Nanoparticle stability studies

The UV-vis absorbance was measured in a Nano-Drop 1000 Thermo Scientific spectrophotometer, with a path length of 0.1 cm. Graphics were obtained using the ad-hoc program ND-1000 V3.71. For FT-IR experiments an FTIR Affinity1-Shimadzu GladiATR was used with a laser lamp with a HAPP-GENZEL absorbance, the range was from 400 to 4000 cm-1, with 120 scans per sample and a resolution of 4 cm-1; the cell was of CeZn with 45° of inclination. For fluorescence studies, an S2 Scinco Fluorspectrophotometer was used.

In the ionic stability and profile release experiment, a MICRO 17 TR (Micro High-Speed Centrifuge) centrifuge was used. In order to measure D.L.S. samples a Zetasizer Nano ZS Malvern Instruments Ltd, Worcs, Malvern, UK was used.

#### 2.4.1 Stability assay at different pH

From stock samples of 450 μM (BSA) and 555 nM (BSA NP), a dilution of 45 μM and 55 nM respectively were used for the assay. Samples were incubated at different pH solution values (2.0; 7.0; 9.0) from buffer PBS following protocols from the CRC Handbook of Chemistry and Physics [21]. After an hour of incubation, Fluorescence and FTIR assays were carried out.

For the FTIR study, 500 μl of each sample were blow-dried in a ZnSe lens with 45 degrees of inclination. The spectrums were recorded with 120 scans at a resolution of 4 cm^−1^. Data obtained was analysed by Software IRSolution. For the fluorescence study, dilutions of the samples were made to a final concentration of 4.5μM (BSA) and 5.5 nM (BSA NP). The λex was 280 nm, and the λem was 337 nm in order to excite the Trp and Tyr in the vehicles. Slits were 2.5 nm opened, and the measurement was carried out at a speed of 60 nm/min. The recording range went from 290 nm to 400 nm. Triplicates of each sample were recorded. For graphics and data analysis the software used was Prism GraphPad v. 5.0.

##### 2.4.2a. SDS denaturation curve

Stock samples of BSA (450 µM), BSA NP (555 nM) and SDS (450 μM), was diluted keeping the vehicle concentration constant at 2.25 µM and 2.77 nM for BSA and BSA NP, respectively. SDS’s concentration varied from 0.00 – 16.00 µM.

These samples were measured by fluorescence; the λex was 280 nm, and the λem was 337 nm (exciting both Trp and Tyr). Slits were 2.5 nm opened, and the measurement was carried out at 60 nm/min. The recording range went from 290 nm to 400 nm. Triplicates of each sample were recorded. The ratio between the vehicle and the detergent was calculated by equation (1)

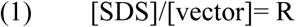

##### 2.4.2b. Urea denaturation curve

Measurement procedures were carried out the same way as described for SDS stability. BSA NP samples were measured against a wide range of increasing urea concentration (0.0-9.0 M) maintaining the nanoparticle and BSA concentration constant at 4.5 μM and BSA NP concentration at 5.5 nM.

##### 2.4.2c. Tween 80 denaturation curve

As regards stability with the chaotropic agent Tween 80, the BSA concentration was of 4.5 μM and BSA NP of (5.5 nM), while the chaotropic agent concentration went from 0% Tween 80 up to 90% v/v. Data obtained was analysed as described for SDS and urea stability assays.

#### 2.4.3. Ionic Strength Stability

Samples of BSA NP (25 nM) and BSA (20 μM) at pH 7.0 were incubated with increasing concentrations of NaCl and KCl separately (0.0 – 1.9 M). Incubation lasted for an hour at 4 °C. Then they were centrifuged for 10 minutes at 4000 g. The supernatant of each sample was measured at 280 nm. Data were analysed with GraphPad Prism 5. Statistical analysis was done using Two-Way ANOVA, post-test Dunnett, and Tukey’s test.

#### 2.4.4. Stability of deviant behaviour through time

BSA NP concentration was constant 555 nM in a 30 mM PBS pH 7.0 solution. Samples were kept at 5 °C during the experiment. Fluorescence, FTIR and D.L.S measurements were made at the beginning and at days 15; 30 and 60 of the experiments. Data were analysed by GraphPad Prism v 5.0

### 2.5. Folic Acid attachment to the BSA NP

The folic acid attachment was carried out as described in Du et al., 2013 [22]. Briefly, a 5 mg/ml folic acid solution (FA) with 3 mg of 1-ethyl3-(3-dimethyl aminopropyl) carbodiimide and 2 mg of N-hydroxy-succinimide was kept under constant stirring in the dark at room temperature. Incubation lasted 4 hours. After, a 30 mg/ml BSA pH 7.0 solution was added for every ml of the FA solution. This solution was kept under dark at room temperature overnight.

The FA-BSA was separated from free FA by centrifugation in an ad hoc column of 1 ml Sephadex G50 at 2000 rpm for 2 minutes at 4 °C. An ethanol BSA solution was added to the FA-BSA solution until reaching 35 % v/v of ethanol in the solution composition. The solution as it was gamma irradiated by a 60Co source (at PISI CNEA-Ezeiza, Argentina), at a dose rate of 1 kGy/h (lowest dose set at 2 kGy and upper limit dose set at 10 kGy). The sample temperature during irradiation was between 5-10 °C. After, samples were eluted in a molecular exclusion column to exchange the ethanol solution for 30 mM PBS pH 7.

In order to estímate the process yield, samples were measured for its UV-visible absorbance at 363 nm. The spectrum represents the FA absorbance maximum, which allowed the quantification of the FA attached to the nanoparticle.

The folic acid attachment was carried out as described here and after the irradiation of the BSA NP. Both products were analysed and compared.

#### 2.5.1. Characterisation of the FA-BSA NP

The folic acid attached BSA NP was characterised as previously described by D.L.S, UV-visible, fluorescence and FT-IR spectroscopy.

#### 2.5.2. Interaction study: FA-BSA NP with Emodin (E)

For this study, 18.4 nM of FA-BSA NP dispersion was used together with a 450 µM emodin (E) ethanol solution. Dilutions on the E solution were made in order to follow an [E]/[FA-BSA NP] ratio range between 0.00 – 12000, where the E highest final concentration used was 90 µM. The determination of the dissociation constant (Kd) was carried out as explained in Sevilla et al., 2007 [11].

#### 2.5.3. A cytotoxicity study in MCF-7 cells treated with FA-BSA NP

The functionality of the FA-BSA NP and the bioconjugate FA-BSA NP/E was evaluated through a metabolic activity assay in the human tumor breast cell line MCF-7. Cells were kept at 37 °C with a 0.5 % CO2 injection with MEM media, 10 % v/v fetal bovine sera.

The assay was carried out through the reduction of tetrazolium dye MTT 3-(4,5-dimethylthiazol-2-yl)-2,5-diphenyltetrazolium bromide to its insoluble formazan form. Cells in a 96 well plate were incubated with 55 nM FA-BSA NP and 45 µM E solutions for 4, 24 and 48 hours (groups of n=12). After incubation, 100 µl of a 1/ 10 of 2 mg/ml MTT solution was added to each well. After 45 minutes incubation, the plate was incubated with 200 µl DMSO/ well, and the well absorbance was measured at 590 nm. Control cells were those with no treatment. Thus they represent 100 % of metabolic activity. Results represent triplicates of the assay.

### 2.6. Cell immune secretion by NPs

The samples studied in this section were increasing concentrations of BSA; BSA NP; lyophilised BSA NP in PBS 30 mM, pH 7.0 and FA-BSA NP.

#### 2.6.1. Murine RAW-BLUE immune response

Cells were cultured at a concentration of 5 e5 cells per well in a 48 well plate, 24 hours before the experiment. Incubation with the samples lasted 22 hours, where samples were diluted in complete media. At the end of each incubation, the supernatant was removed and preserved for future experiments. An MTT assay was carried out on the remaining cells. LPS 100 µg/ml served as positive control, while cells incubated in complete media without stimulant served as negative control.

#### 2.6.2. Cell Metabolic Activity Essay

Briefly, after incubation with the stimulants, 150 µl of 0.2 mg/ml MTT was added to each cell with cells. After 2 hours of incubation at 37 °C, 5 % CO2, the plate was washed, and 150 µl of DMSO was added per well. The plate was read in a BioTek plate reader at 570 nm.

#### 2.6.3. NFκB transcriptor activation

A total of 20 µl of the collected samples were placed in a 96 well plate. A total of 180 µl of Quanti – Blue was added to each well-containing samples. Incubation lasted for an hour at 37°C, in the dark. Signal was read using a BioTek plate reader at 620 nm. Data were normalised to the value of cells without any stimulant and then to the MTT data collected previously. Data was represented by Software Excel v. 2015 and GraphPad Prism v 5.0.

#### 2.6.4. Cytokine expression

In a 96 well plate for ELISA a total of 30 µl of the collected supernatant were used per well. Briefly, the ELISA plate was prepared with 1 % BSA solution and coated with the capture antibody. Then, the plate was incubated for 2 hours with the standard and samples solution. After the incubation, the plate was washed and first, the detection antibody was incubated and then, the substrate. Finally, the stop solution awarded each sample with colour depending on the cytokine concentration in it. The plate was read in a BioTek plate reader set at 450 nm.

## 3. Results and Discussion

### 3.1 Size exclusion chromatography and SDS gel electrophoresis

Nanoparticles formed from BSA molecules (BSA NP) due to desolvation in an ethanol solution were then stabilised by γ irradiation [3][4][5] (Figure 1a). After the preparation process, the NPs were separated by a size exclusion chromatography column (Sephadex G – 200) into different populations according to the order of elution; those of a larger size eluted first, followed by the smaller population (Figure 1b). The aliquots were then identified by UV-Vis spectroscopy: the first maximum was identified as NP, while the second maximum was identified as residuary albumin. With the aid of the elution profile, the yield of the NP was obtained; the colloidal solution contained 80 % NP and 20 % of residuary free BSA molecules (Figure 1b).

**Figure 1.**
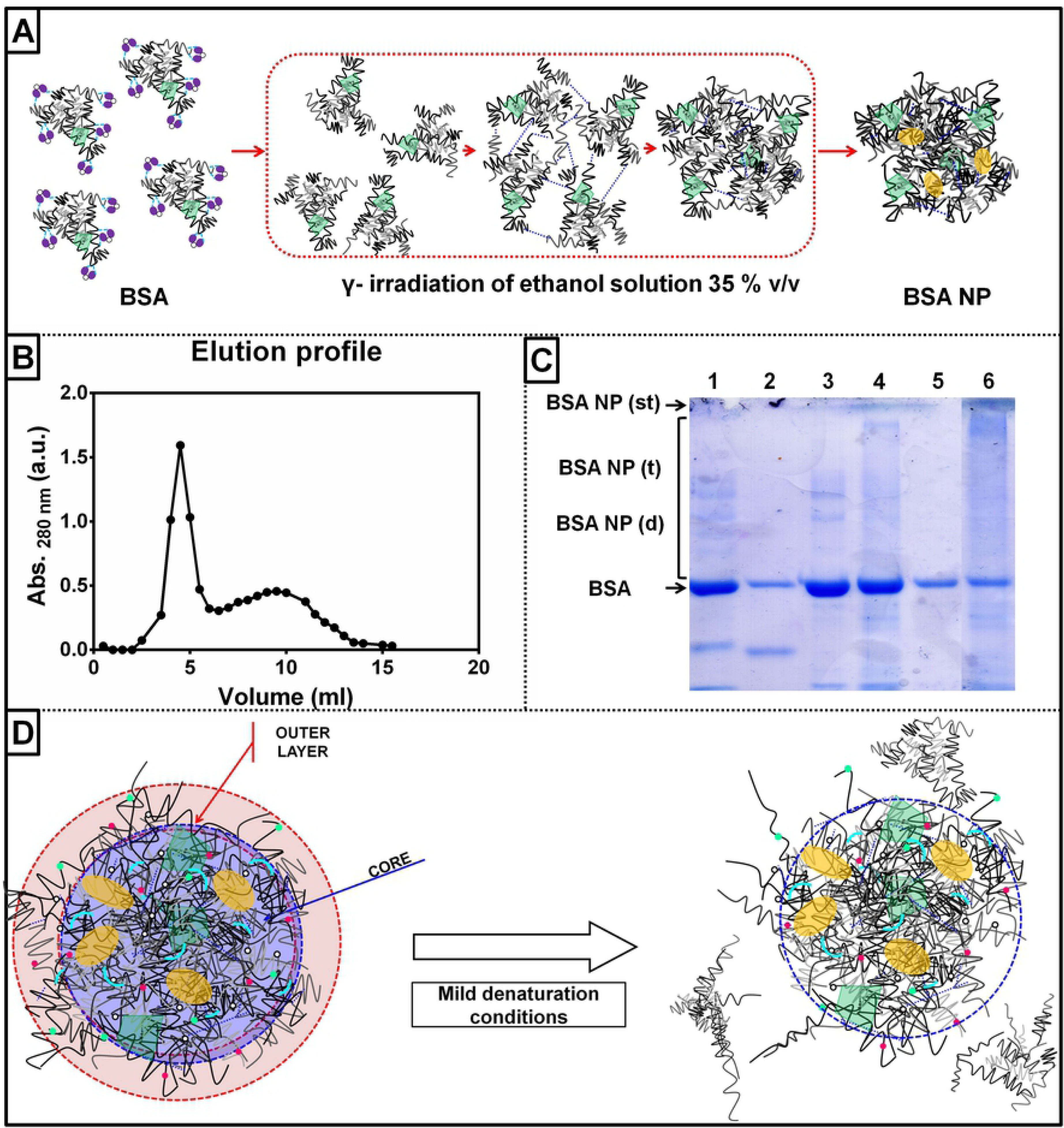
(a) Illustration of the process of irradiation by which the BSA NP is obtained. (b) Elution profile obtained from a size exclusion column of the BSA NP. (c) SDS-PAGE of BSA NP samples; 1) protein molecular markers (BSA trimmers, dimmers and monomers), 2) protein molecular markers (BSA, ovalbumin, lysozyme), 3) BSA in buffer PBS 30 mM, pH 7.0, 4) BSA NP in ethanol solution 30 % v/v, 5) BSA NP in buffer PBS 30 mM, pH 7.0, 6) BSA NP after separation in a G200 Sephadex size exclusion chromatography column. The bands corresponding to BSA, BSA trimers (BSA (t)) and dimers (BSA (d)) are indicated in the Figure, together with the nanoparticle that stayed in the stacking gel (BSA NP). (d) Illustration of the BSA NP outer layer and core with different forces involved in each section according to results in Figure 1 (c).

An SDS-PAGE gel study took place in order to study the BSA NP assembly (Figure 1c). If weakly linked proteins form the NP, the mild denaturation experimental conditions in which the gel is carried out (SDS presence in gel) would be enough to disaggregate the BSA NP. Contrary to this, if strong forces crosslink the BSAs, then the NP will not run, offering a distinguishable one spot protein aggregate in the stacking gel. Neither condition is desirable; the BSA NP should have stability balanced between both extremes, as it must preserve its function as a carrier and biodegradability after delivering the drug.

The assembly of the NP was tested in different conditions (Figure 1c). The samples were: BSA in phosphate buffer 30 mM PBS, pH 7.0 (lane N°3); BSA NP in ethanol solution (30 % v/v) as obtained after γ irradiation (lane N° 4); BSA NP in buffer 30 mM PBS, pH 7.0, after size exclusion column (lane N° 5); and after separation through a molecular exchange chromatography column (Sephadex G 200) (lane N°6). Two different protein markers were used: one containing BSA, ovalbumin and lysozyme (lane N° 2); and another one of our own making containing BSA trimers, dimers, and monomers (lane N° 1). It was possible to observe different aggregate sizes in each sample. All gel lanes showed different concentrations of free BSA (66 kDa). The stacking gel showed stains corresponding to NP that did not run in every lane. BSA NP Sephadex G200 (Figure 1c, lane N° 6), was the sample with the highest concentration of NP; 68 % of NP distributed; 48 % in the stacking gel and 20 % of different NP sizes. This NP was also the one with the least concentration of free BSA (31 %). The rest of the BSA NP samples showed a higher concentration of free BSA, (30 - 80 %) and an NP concentration ranging from 8 to 30 % (Figure 1c). Because of the profile of the running gel, it is suggested that different forces assemble the BSA NPs. When subjected to mild adverse experimental conditions, like an SDS-PAGE, the NP would lose some of the attached BSA molecules in its composition. The lost molecules represent the weakly external linked layer (Figure 1d). As none of the BSA NP samples, except for BSA NP Sephadex G200, was not separated from the residuary protein after irradiation, they showed a large percentage of free BSA (Figure 1c). As a result, the weakly linked BSA molecules in the NP joined the residuary BSA in the sample (20%), resulting in a higher percentage of free BSA (over 20%). This weakly linked protein layer could be described as an external layer protecting a deeper albumin core where the protein molecules were crosslinked by stronger forces that are difficult to denature. The different bands observed in every lane seeded with BSA NP are also indicative of the presence of different NP sizes resulting from the NP degradation (Figure 1d).

### 3.2 Structure, morphology and size

The size of the NP obtained was of 70 nm of diameter and a Z potential value of −25 mV confirming data previously reported [5]. To further characterise the BSA NP, different microscopy techniques were used to observe the shape of the aggregates. According to T.E.M, the BSA NP has a spherical-like shape with a given mean diameter of 35 nm (Figure 2a-b). The results represent a monomodal population with a minor polydispersity (35.4 ± 0.35 nm) (Figure 2c). The microscopy also shows aggregation of the sample (Figure 2a-b). Nevertheless, this phenomenon could be because of the dehydration the sample undergoes during experimental preparation.

**Figure 2.**
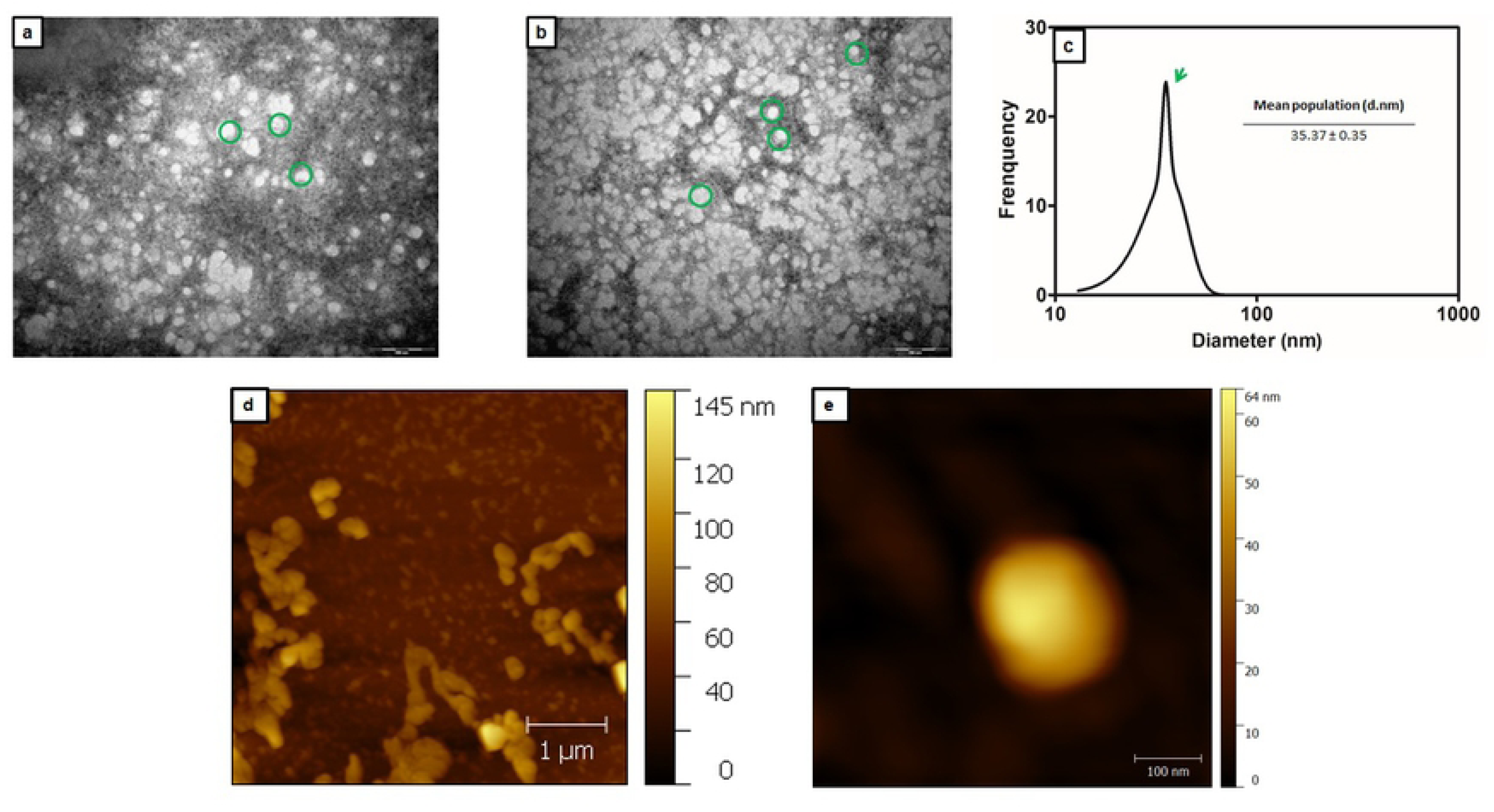
Transmission electron microscopy (T.E.M) of the BSA NP (a-b). Microscopies represent different samples fields were the magnification correspond to 50.000 X, were 1 cm= 200 nm. 20 microscopies were taken, and a total of 450 NP were counted. (c) T.E.M histogram represents the mean size of the total of NP analysed. Atomic Force Microscopy (A.F.M.) of the BSA NP (d-e) Microscopies represents the aggregation in the sample while dehydrated state for A.F.M. (d). Magnified single BSA NP (e).

The A.F.M showed aggregation of the BSA NP while in the dehydrated state, together with BSA molecules in the background (Figure 2d). A magnification of a single BSA NP particle showed defined edges and heterogeneous surface (Figure 2e). The height of the NP obtained in the microscopy corresponds to 64 nm, which is within the normal measurements for the BSA NP obtained by D.L.S measurements [5].

As stated before, according to the TEM taken, the NP’s diameter resulted in approximately 35 nm, which coincides with our previous results in ethanol buffer solution in the range 20 - 40 nm [3]. Differences among the NP sizes obtained using different techniques might be explained as follows: BSA NP for TEM and BSA NP in ethanol solution provide a dehydrated state; in a buffer solution, the NP is hydrated and therefore its diameter varies into 70 nm. According to these parameters, each NP is formed by a total of 2246 BSA molecules. The diameter of the NP implies that as a carrier, the ratio between volume and surface would be optimised; and thus, the surface tension will be reduced. Its shape also allows a better circulation if IV administered; therefore, the NP will be able to avoid attachment to the vessels [16].

AFM showed that the BSA NP has an ellipsoidal shape and irregular surface. The albumin does not have a uniform surface, alongside, during the NP preparation process, a little denaturation and structure alteration on the albumin’s part is typical due to the steps involved in the preparation process (ethanol desolvation and gamma irradiation). Therefore, the irregular surface feature of the NP is corroborated (Figure 2d-e).

### 3.3. Spectroscopy methods

Spectroscopy experiments were carried out in order to further characterise the BSA NP. Detecting any alteration in the BSA molecules forming the NP might imply an overall loss of function. The preservation of function in the BSA NP is essential as it is aimed at improving drug delivery in medical treatments.

Unlike albumin UV-vis spectra (shoulder at 283 nm), the UV-Vis profile of BSA NP shows the Trp_BSA NP_ shoulder at 277 nm and a shoulder broadening of 7 nm (from 18 to 25 nm)(Figure 3a). According to Rohiwal et al., 2016 [23] a broadening of the shoulder in the BSA NP spectra means not only conformational alterations in the structure of the proteins forming the NP but an alteration in size of the particle in the colloidal suspension (Figure 3a). Further spectroscopic experiments were carried out in order to elucidate these alterations in structure.

**Figure 3.**
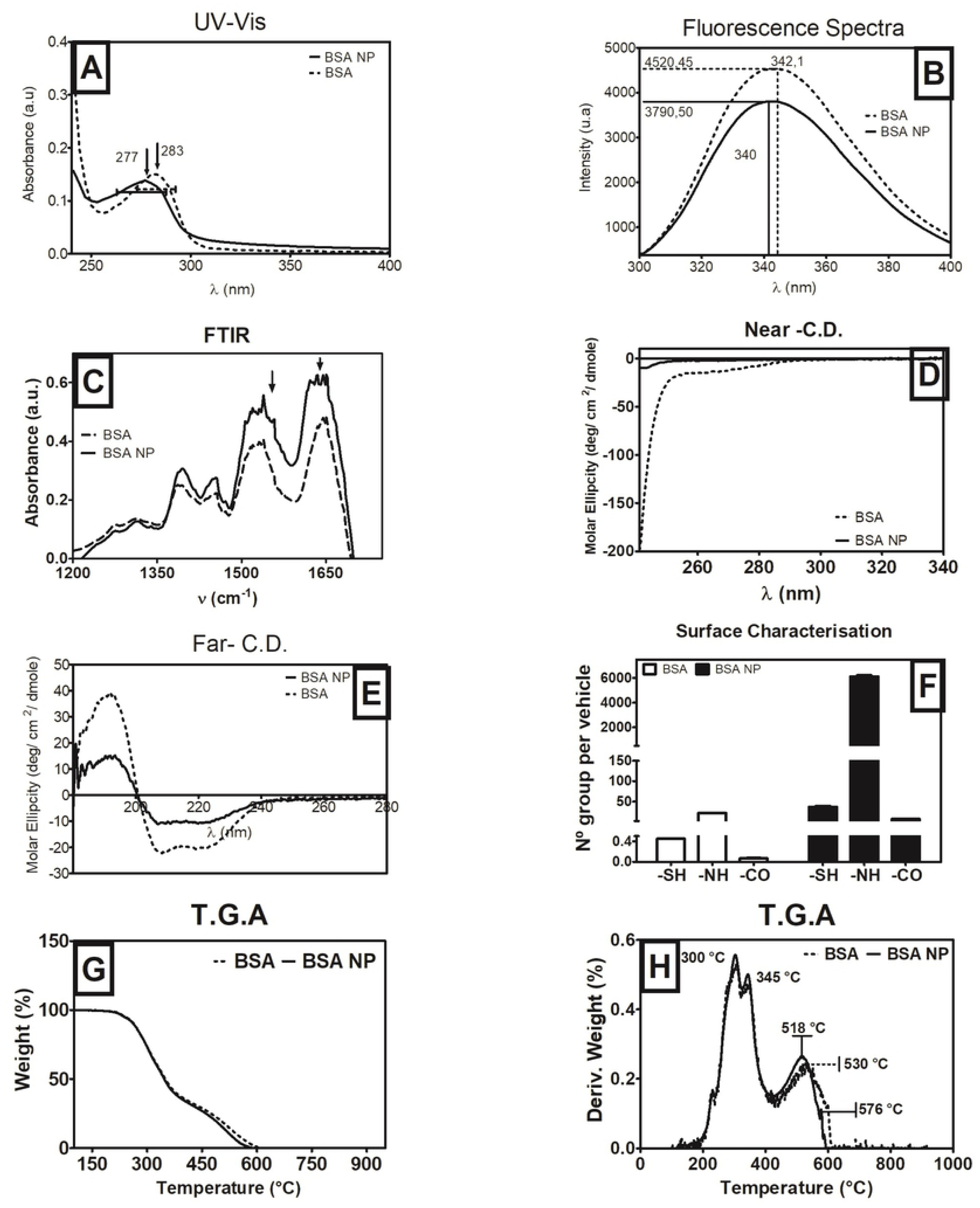
Biophysical studies carried out on the BSA NP; (a) UV-vis spectroscopy profile of BSA (dotted line) and BSA NP (full line) _ arrows indicate shoulderś maxima. (b) FT-IR study of BSA (dotted line) and BSA NP (full line) _ arrows indicate α-helix maximum and β-sheet-like structures signal (80 scans with a resolution of 1.00 cm^−1^). (c) Fluorescence study of BSA (dotted line) and BSA NP (full line) _ arrows indicate where the maximum intensity of Trp is set (λex= 295 nm and λem= 377nm; slits: 1.5 nm). C.D. spectroscopy profiles of BSA NP near UV with signals corresponding to aromatics amino acids (Trp, Tyr, and Phe) (d), and far UV with signals corresponding to α-helix and β-sheet structures. Experiments correspond to triplicates (e). The surface characterisation for – SH, −NH and –CO groups exposed to the BSA NP surface (f). TGA profile of BSA (dot line) and BSA NP (full line) from 100 – 920 ªC (g). The first derivative of BSA (dot line) and BSA NP (full line) from the data in (g): the central maximum in the first derivative profile are signaled in the figure (h). The data described correspond to duplicates.

The BSA NP FT-IR spectra showed that the 1652 cm^−1^ maximum representing the α-helix composition of the protein, did not experience any displacement when the NP was formed (Figure 3b). This maximum is of great importance as it is sensible to the environment and any change in the protein structure. Hence, the alpha - helix content, the most critical domain in the secondary structure of the protein, is not lost. A band signal corresponding to β-sheet structure signal (1510 – 1545 cm^−1^) is described for the NP profile, whereas this maximum is absent in the BSA profile (Figure 3b). While forming the NP, the dehydration of the BSA molecule is significant enough to bring the molecules together in an attempt to compensate for the H-bonding loss in the absence of water due to the desolvation process. These bonds are believed to help maintain the β-sheet structure: the BSA NP spectra showed less CO, CN and NH groups’ freedom degrees (groups prone candidates to this kind of structures). While Rohiwal et al., 2016 [23] obtained more significant blue shift of the amides I, II and III, their BSA NP was prepared by desolvation process with a crosslinking method in the presence of glutaraldehyde. The changes observed in Rohiwal et al., 2016 [23] work are observed as frequency displacement to lower frequencies, confirming that structural stability has been changed [23]. The difference of the crosslinking method used might be reason enough for the differences in the spectra obtained. The fact that there is also a frequency displacement of the carbonyl groups that do not accept H bonding, from 1600 cm ^−1^ to 1555 cm^−1^, means that the NP has a more compact structure than BSA [24].

By fluorescence spectroscopy, it was observed that the BSA NP fluorescence profile showed a slight displacement of 2.5 nm to lower wavelength (blue shift) meaning a more hydrophobic Trp_BSA NP_ environment confirming UV-visible data (Figure 3c), whereas, if the BSA NP was obtained by glutaraldehyde desolvation method, it experiences a significant blue shift of 16 nm [23]. Hence, by using γ irradiation methodology, the BSA structure is more preserved in the NP. Moreover, the spectra showed a slight emission decrease in the Trp_BSA NP_, describing a possible quenching effect among the various Trps present in the BSA NP due to molecular crowding forming the NP [5]; the Trps (143 and 212) are closer to each other, generating a quenching effect (Figure 3c). It could be inferred from the above that the albumin forming the BSA NP has a more compact conformation than free BSA.

The C.D. spectra from BSA NP showed aggregation of the molecules forming the NP in the secondary structure in both near and far CD (Figure 3d-e). The aromatic signals in the near – CD gave a low signal in the BSA NP spectrum (Figure 3d). The same was observed in the α-helix and β-sheet signals in the far – CD (Figure 3e). Both results are indicative of structural alteration in the BSA molecules such as protein aggregation.

The methodologies used as γ irradiation are expected to generate surface modifications of the functional groups present in the surface of the NP due to ROS formation. Detecting the presence of these groups is useful for describing how the BSA molecules are bonded between each other, or any other alteration structure. Experiments were carried out to detect how many –SH, −NH and –CO superficial groups the BSA NP has.

Results showed considerable amount of all functional surface groups present in the NP (Figure 3f). According to the increased number of functional groups in BSA NP, the differences in –SH and −NH mean that irradiation brakes and rearranges the –S-S binding groups and possibly the origin for the protein aggregation (Figure 3f). BSA is known to have one –SH group per molecule, when forming the NP the final –SH superficial groups per molecule are 37. This result states aggregation and reorganization on the structure of the protein molecules forming the NP as it is composed by a total of 2246 BSA molecules [5]. The non-detected –SH might be forming intermolecular S-S stabilising the NP.

Moreover, taking into account the number of NH per BSA (30) and the number of NH detected per nanoparticle (6122), 9% of the total amount of –NH groups (67380) are exposed to the surface (Figure 3f). According to this result, there will be approximately 200 BSA molecules exposed to the surface of the NP, while the rest of the –NH might be involved in the crosslinking of the BSA of the NP. These groups might be part of the β-sheet-like structures described in the FTIR results (Figure 3b). The increased amount of superficial −NH will ease the covalent binding of a ligand to the NP. Even though the presence of superficial CO is related to protein oxidation [25], it is also a sign of structure alteration due to the irradiation process experimented by the BSA molecules. The CO also can be forming the β-sheet-like structures described in the FTIR results together with the –NH groups. Therefore, the presence of these groups might be desirable in order to stabilise the aggregate into a BSA NP.

Data collected from spectroscopy experiments showed changes in the UV-Vis absorbance profile of the BSA NP compared to the BSA spectrum. Not all of the albumin molecules forming the NP preserve its structure after protein aggregation (Figure 3a). From the C.D. experiment, protein aggregation was observed in the BSA NP spectrum. Thus, might be because when forming the NP, the desolvation process with ethanol and γ irradiation process promotes protein aggregation (Figure 3e-f). Results in the FT-IR and fluorescence experiments aid the aggregation theory built to explain the signal loss observed in the C.D. experiment (Figure 3b-c).

### 3.4 TGA measurements

It was observed by TGA measurements differences between the BSA and BSA NP degradation profile (Figure 3). The early drop of the curve signals the NP as a structure more sensitive to heat at higher temperatures; the sudden weight loss after 300 °C in both samples is due to the loss of molecules such as carbon dioxide and ammonia [26] (Figure 3g). The derivation of each curve states that BSA NP and BSA had a similar behaviour at temperatures lower than 450 °C, even though there is no displacement of maximum at 295 and 310 °C in the NP profile in comparison with the BSA profile but values reached were slightly higher in the case of NP than BSA. It can be observed that BSA NP reached after 500 °C a maximum 12 degrees before BSA and the earlier drop near 0 % of remaining weight (Figure 3h). It is possible that the shift of the maxima at 518 °C in BSA NP and 530 °C in BSA was a consequence of preparation process of nanoparticles, proteins were slightly affected by irradiation in bonds and conformation as can be seen partially in FT-IR section. This difference is of significance indicates the NP is composed by interactions more prone to be destabilised by heat than BSA.

Differences in thermal degradation between NP and BSA were also found by other authors who reported a shift to lower temperatures in thermal degradation, as a sign of protein organisation due to new bonds between albumin molecules [27]. Singh et al. 2017 [28] have reported that degradation (pyrolysis, in nitrogen) became earlier in nanoparticles made with BSA concerning control BSA in the range of 250-450 °C. The reorganisation in structure present in the BSA molecules after the NP preparation process was not only observed by FT-IR, but also by surface characterisation studies. Both suggested the existence of BSA molecule bonding by β-sheet-like structures (FT-IR), and the possibility of S-S intermolecular bonds (Figure 3c; 3f). As it is, the T.G.A study confirms the existence of such bonding between albumin molecules during the BSA NP preparation process.

In order to study the stability of the BSA NP, it is necessary to understand that different forces play a specific role in preserving NP stability. The BSA NP AFM microscopy (Figure 2e) describes it as a sphere-shaped, with a non-uniform surface formed by moieties where proteins are protruding and moieties of dark areas where proteins are embedded forming the core of the NP. These results suggest different attractive and repulsive van der Waals interactions playing their part in the intermolecular bonding between albumin molecules, as well as covalent union prevailing at the NP core level.

Depending on the solution the BSA NP is in, are the forces acting upon the protein interaction. In a saturated solution, the attractive interaction of the proteins in the NP is solely provided by short-range hydrophobic interaction [29][30][31]. As this type of interaction happens at a length scale of less than 1 nm, it can be assumed that the spatial scale is given by the water layer (c.a. 0.2 nm) depending on the polarity of the solution [32][33][34]. On the other hand, the repulsive interaction is given by the electrostatic interaction between the nucleus protein aggregate and the addition of protein monomer or other nucleus when in the making process, varying from the lowest to the highest ionic strength solution reported (c.a. 0.8 – 2.5 nm) [33][34][36]. The stability of the NP was analysed according to the alteration of the structure depending on the moieties exposed.

### 3.5. NP-Stability against pH

The importance of the stability of the BSA NP relays on the inoculation root as a nanovehicle for either oral or IV administration. Drug delivery IV or oral administration means that the nanovehicle will be exposed to different pH media. If administered orally, the NP should resist denaturation at acidic pH values. If administered IV, the NP should be stable at physiological pH value. Besides, considering that different dispersion procedures may lead to variable results the behaviour of the NP also needs to be tested in different pH solutions. Thus, the BSA NP stability in different pH media will help determine their behaviour in this environment and to acquire fundamental understanding in order to determine how the physicochemical properties influence biological protein nanoparticle structure/function.

As it was, three pH values were tested: 2.0, 7.0 and 9.0. The acidic pH value 2.0 corresponds to that of gastric juice, whereas pH 7.0 represented a solution near physiological value where the protein of the NP is in its native form. The pH 9.0 solution was chosen considering the BSA conformation change from N to B in an alkaline pH [36].

FT – IR experiment shows similarity in the behaviour of the NP with that of the BSA in its molecular form (Figure 4.1a-c). To facilitate data analyses comprehension: signals in Amide I correspond to the secondary structure of the protein: at 1500 cm^−1^ there is a β-sheet signal, and at 1650 cm^−1^ α-helix vibrations can be observed at pH 7.0 (Figure 4.1a). In the same spectrum, between 1300 and 1450 cm^−1^, Amide II, signals from C=C, C=O, C-C appear. The spectrum profile acquired at pH 7.0 was used as a reference against the ones acquired from the other two solutions.

**Figure 4.**
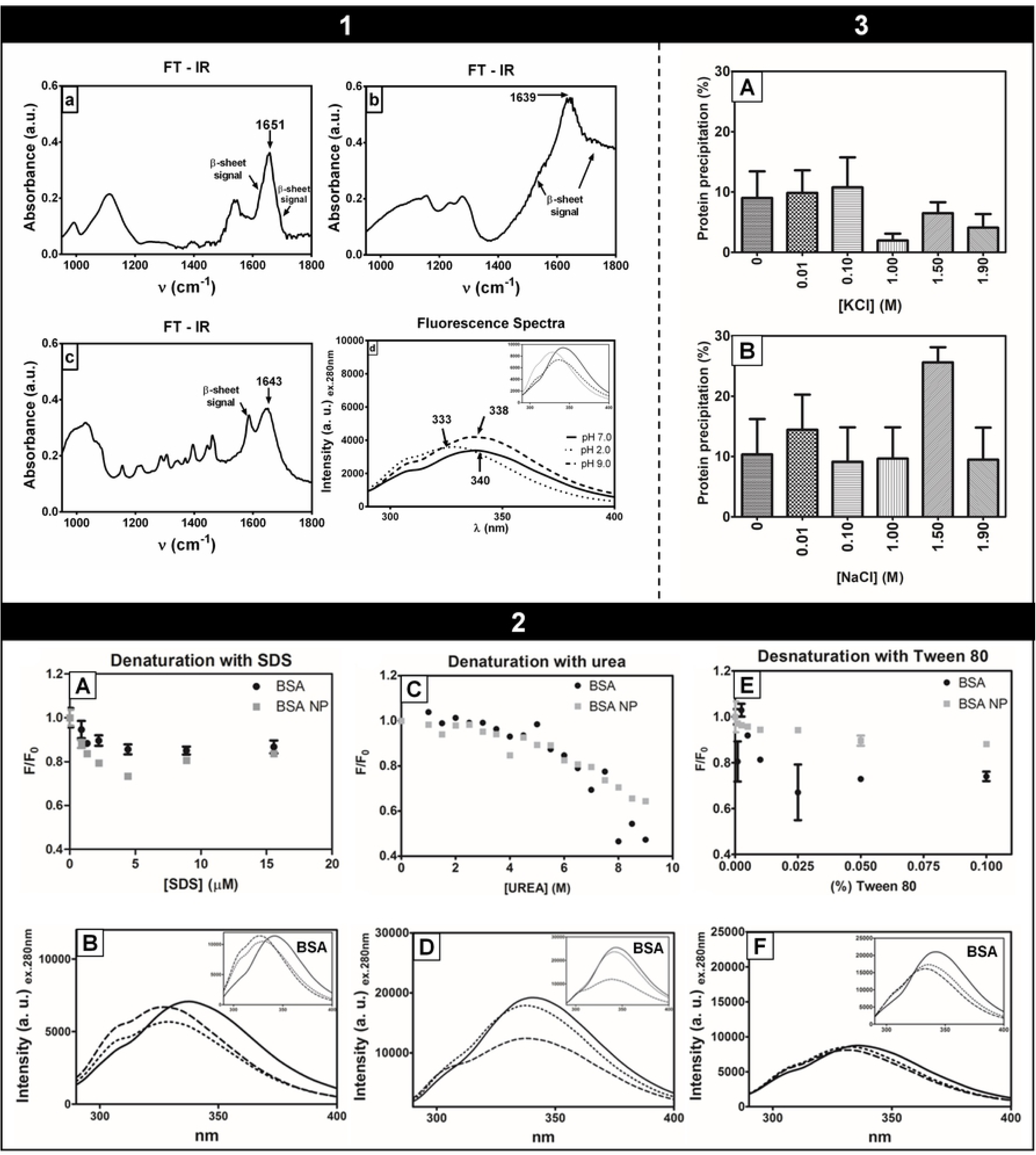
1 BSA NP spectra obtained by FT-IR spectroscopy (a-c). FT-IR measures were of 100 scans at a 1 cm-1 resolution. (a) Samples at pH 7.0; (b) samples at pH 2.0; (c) samples at pH 9.0. Fluorescence measures were triplicated. (d) Fluorescence spectroscopy of BSA NP at different pH values with inset of the BSA fluorescence spectroscopy of BSA. 2. Behaviour of the BSA NP under the influence of different chaotropic agents. (a-b) SDS: (a) denaturation curve using increasing concentrations of SDS where [SDS]/ [BSA NP]= 0.0 - 5617; (b) fluorescence emission spectra of the three species of the BSA NP behaviour with increasing concentrations of SDS, with inset of the BSA behaviour treated with increasing concentrations of SDS. (c-d) Urea: (c) denaturation curve using increasing concentrations of urea where [urea]/ [BSA NP]= 0.0 - 36; (d) fluorescence emission spectra of the three species of the BSA NP behaviour with increasing concentrations of urea, with inset of the BSA behaviour treated with increasing concentrations of urea. (e-f) Tween 80: (e) denaturation curve using increasing concentrations of urea where [Tween 80]/ [BSA NP]= 0.0 - 3249; (d) fluorescence emission spectra of the three species of the BSA NP behaviour with increasing concentrations of Tween 80, with inset of the BSA behaviour treated with increasing concentrations of Tween 80. 3. Protein precipitation using different salts NaCl (a) and KCl (b) for BSA NP. Data were analysed with ONE-WAY ANOVA, post-test Dunnett P<0.05 using BSA NP as a control. Measurements were triplicated.

At pH 2.0, (Figure 4.1b) signals from C=C, C=O, and C-C are present, but at lower frequencies (blue shift): the α-helix signal shifts from 1651 to 1639 cm^−1^ (c.a. 12 cm^−1^). The blue shift of the α-helix peak in Amide I indicates that the groups involved have loosened from their interaction with each other, allowing to move more freely, indicative of a change in the secondary structure.

The spectrum at pH 9.0 describes signals for the regions of interest at lower frequencies (Figure 4.1c). These signals present frequency displacement of several degrees, indicating less α-helix structure. The α-helix signal shows a smaller blue shift than that at pH 2.0, from 1651 to 1643 cm^−1^ (c.a. 8 cm^−1^). However, as the alteration of the spectra at pH 9.0 is not as pronounced as the one at pH 2.0, the structure-loss experienced by the NP at the alkaline pH is different from the one experienced at acidic pH.

The fluorescence emission of the Trp in the main hydrophobic binding pocket site of the BSA is usually used to study the behaviour of changes in the surrounding groups. With the aid of this study, an estimation of the hydrophobic character and protein denaturation grade can be asserted. The same reasoning can be extended to the Trp of the BSA NP.

The fluorescence emission profile acquired at pH 7.0 was used as a reference for future comparison with the different pH values tested (Figure 4.1d); the maximum peak is representing the TrpBSA NP emission considering native experimental conditions is set at 340 nm. At low pH values (2.0) a blue shift from 340 nm to 333 nm, whereas, at pH 9.0 a blue shift is reported from 340 nm to 338 nm (Figure 4.1d). Both changes indicate a more hydrophobic microenvironment surrounding the TrpBSA NP. Itri et al., 2004 [37], correlates an increase in the fluorescence emission with greater exposure of the Trp to the surface. Considering this, at pH, 2.0 and 9.0, the TrpBSA NP is nearer to the solvent compared to its position when the NP is in a native conformation (pH 7.0). Both show structure and group alteration of the BSA NP at these extreme pH values.

There are many studies regarding the BSA behaviour under different conditions, and its structure changes undergo in each condition. For example, Michnik et al., 2005 [36], described by thermal denaturation different states that BSA goes through while changing the solution’s pH value. Between pH 4.5-7.0, BSA adopts a standard “N” form. In the range of pH 4.0-4.5, the transition towards a fast migrating “F” form occurs. The transition is completed abruptly under pH 4.0. From pH 3.9 onwards the form predominating is “E,” expanded. Here, the BSA is completely depleted from its secondary structure. The opening of the secondary structure begins with the “N” -> “F” transition [36]. According to Barbosa et al., 2010 [38], there is another transition taking place between pH 8.0 and 9.0, which is named as basic transition “B”. In this case, the protein loses some of its rigidity. Thus its ratio increases slightly.

Based on previous works and our findings, similar behaviour is proposed for the BSA NP under different pH values. When in acidic conditions, the BSA structure presents a fully extended structure, denoting maxima dimension and asymmetry [38][39]. The NP might present what it is to be considered an extended version of it. This change is given by a relaxed form with a tendency to a loose-disaggregation of the covalent bound albumin molecules in the NP (Figure 1e). As the proteins in the aggregate presented structure conformation changes, meaning some of its hydrophobic moieties are exposed [5]. This conformation is comparable to the molecular albumin denatured E form. The NP form does not fit with that of the NP denatured state equal to E form, but has its particular conformation where its structure shows great denaturation. Consistent with our assumptions, Li et al., 2016 [40], described a higher concentration of β-sheet structures and a decrease in α-helix content in the BSA at pH values under 3.0. The loss of in α-helix content in the molecular BSA was observed here as a displacement to lower frequencies, whereas, the acquisition of β-sheet content was observed as a signal appearance in the range between 1500 and 1600 cm^−1^. At pH 2.0 the displacement of the α-helix signal to lower frequencies (c.a. 12 cm^−1^) indicates a loss of secondary structure in the NP (Table 1). There is also the appearance of β-sheet signals (Figure 4.1b). Therefore, the NP under this condition would present its own relax NP “E” form (Figure 4.1e).

**Table 1.**
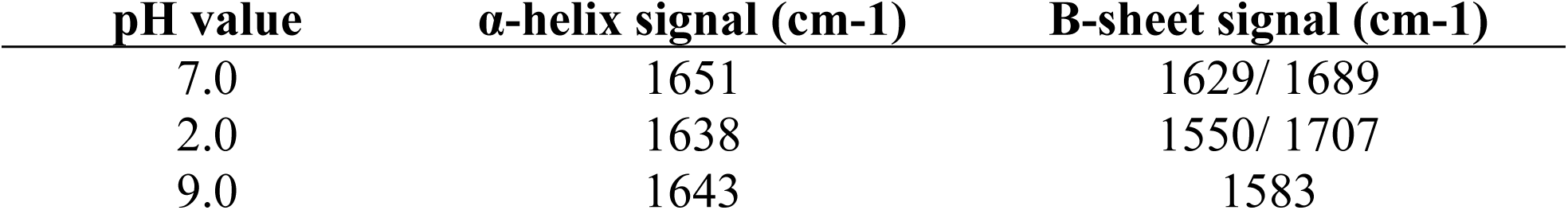
Maxima of the main peaks observed in the FT-IR spectra at different pH values.

At pH 9.0 the NP presents a different degree of conformation changes. Both, loss of α-helix content and gain of β-sheet structures are registered (Figure 4.1c). Probably the greater signal for β-sheet structure indicates more aggregation [36]. Nevertheless, compared with the spectra at pH 2.0, there is less displacement of the α-helix signal (c.a. 8 cm^−1^) (Figure 4.1c) (Table 1). According to these results, it is assumed that the NP changed towards an extended version and halfway aggregated. Therefore, the NP under this condition would present its own NP “B” form: not completely aggregated, but not wholly extended neither (Figure 4.1e).

**Table 1.** Quantification of the ligand molecules per NP based on experimental data from Figure 1.1b-c.

| [BSA NP]<br>(mg/ml) | MW (Da) | NP<br>molecules/ml | [Ligand]<br>(mg/ml) | MW (Da) | Ligand<br>molecules/ml | Ligand/<br>BSA NP |
| --- | --- | --- | --- | --- | --- | --- |
| 30 | 1,48E+08 | 1,22E+20 | 6,00E-01 | 4,41E+02 | 8,19E+20 | 6,71E+00 |
| 30 | 1,48E+08 | 1,22E+20 | 2,50E-02 | 3,89E+02 | 3,87E+19 | 3,17E-01 |

Fluorescence results are by those above. Gelamo et al., 2002 [10], states that there is a correlation between the hydrophobic character of the Trp vicinity and the wavelength displacement that might take place: at displacement to lower wavelength, more hydrophobic character of the Trp surroundings.

At pH 2.0 the displacement of the BSA NP Trp to lower wavelength is of seven nm, whereas at pH 9.0 the displacement of the BSA NP Trp to lower wavelength is of two nm (Figure 4.1d). The latter shows a small change in the TrpBSA NP environment, which suggests what was priory assumed: the NP at pH 9.0 is in a B form. Contrary to this, the wavelength displacement experienced at pH 2.0 suggests a higher-level of the NP change, suggesting the NP “E” form adoption under these conditions (Figure 4.1e). If loss of structure means loss of functionality, the BSA NP is not useful for orally administrated treatments, due to the risk of loss of efficiency caused by the BSA NP alteration structure at acid pH.

### 3.6 NP Stability against Chaotropic Agents

The stability of the BSA NP was tested in different adverse experimental conditions using chaotropic agents in order to test whether the nanovehicle could be used to protect drugs dissolved in other solvents than water. The selection of chaotropic agents used: SDS, urea, and Tween – 80. The focus was on changes that would help describe the presence of three different species present along the experiment. These species were (1) the nanovehicle in a native form; (2) the complex nanovehicle – chaotropic agent; and (3) the nanovehicle in its maxima conformational changes form [10].

Displacement on the wavelength of the spectra meant changes on the microenvironment of the Trp_BSA NP_ of the binding pocket, increasing or decreasing its hydrophobic character (blue and redshift, respectively). An increase or decrease on the fluorescence intensity meant different depth of hydrophobic environment exposure of the BSA NP Trp to the surface [10][37][38].

#### 3.6.1 NP-SDS denaturation curve

As the concentration of SDS increased, the Trp_BSA NP_ maximum experienced a blue shift, from nm_0_= 335 nm to nm_f_= 325 nm, together with a final quenching fluorescence of 6 % of the initial fluorescence emission (Figure 4.2a-b). When the [SDS]/ [BSA NP] is 1603, the fluorescence spectra shifts from 335 nm to 327 nm and the decrease fluorescence emission is of 20 % of the initial fluorescence. These changes state a more hydrophobic character of the microenvironment surrounding the Trp_BSA NP_ together with lesser exposure of the amino acid to the solvent altering the quantum yield (Figure 4.2b). The endpoint of the SDS denaturation curve ([SDS]/ [BSA NP] = 5617), is represented by a blue shift from 335 nm to 325 nm and a decrease in the fluorescence emission of 6 %. The Trp_BSA NP_ is more exposed than in the previous step described but has an increased hydrophobicity in its vicinity compared to the starting point of the experiment (Figure 4.2b).

The kinetics of the BSA NP structure alteration in an SDS solution is as follows: as the SDS concentration increases alteration in the structure of the BSA NP takes place, enhancing the hydrophobic character of the Trp_BSA NP_ binding pocket while rearranging the amino acid exposure to the solvent (Figure 4.2a). The last structure re-arrangement takes place in two steps: first, the exposure decreases (when the ratio is 1603) but by the end of the experiment, the nano vehicle’s groups are rearranged in order to expose further away from the Trp_BSA NP_ from the solvent.

#### 3.6.2 NP Urea denaturation curve

As the concentration of urea increases, the Trp_BSA NP_ maximum suffers a shift towards lower wavelength, a blue shift, from nm_0_= 340 nm to nm_f_= 337 nm, together with a final quenching fluorescence of 35 % of the initial fluorescence emission (Figure 2c-d). When the [urea]/ [BSA NP] is 7.2 e13, the fluorescence spectra shifts from 340 nm to 336 nm and the decrease fluorescence emission is of 8 % of the initial fluorescence. These changes state an almost similar hydrophobic character of the microenvironment surrounding the Trp_BSA NP_ together with slightly less exposure of the amino acid to the solvent altering the quantum yield (Figure 4.2c). Nonetheless, these changes are not pronounced enough to indicate conformational changes and re-arrangement of the BSA NP. The endpoint of the urea denaturation curve ([urea]/ [BSA NP] = 1.6 e14), is represented by a blue shift from 340 nm to 337 nm and a higher decrease in the fluorescence emission of 35 %. The Trp_BSA NP_ is much less exposed than in the previous step described but has not changed its hydrophobic character surrounding the Trp (Figure 4.2c). Even though the microenvironment surrounding the Trp_BSA NP_ did not change its hydrophobicity, the groups surrounding the binding pocket might experience position rearrangements enabling the Trp_BSA NP_ to be further from the nano vehicle’s surface.

The BSA NP in urea solution undergoes structural changes as the urea concentration increased: the structure of the BSA NP not only changes the hydrophobic character of the Trp_BSA NP_ binding pocket but rearrange the amino acid exposure to the solvent (Figure 4.2b). The exposure of the Trp_BSA NP_ gradually decreases as the experiment develops.

#### 3.6.3 NP-Tween 80 denaturation curve

As the concentration of Tween 80 increases, the Trp_BSA NP_ maximum suffers a shift towards lower wavelength, a blue shift, from nm_0_= 335 nm to nm_f_= 330 nm, together with a final quenching fluorescence of 8 % of the initial fluorescence emission (Figure 4.2e-f). When the [Tween 80]/ [BSA NP] is 1.8 e6, the fluorescence spectra shifts from 335 nm to 332 nm and the decrease fluorescence emission is of 3 % of the initial fluorescence. These changes show changes neither in the hydrophobic character of the microenvironment surrounding the Trp_BSA NP_ nor in the exposure of the amino acid to the solvent (Figure 4.2f). The endpoint of the Tween 80 denaturation curve ([Tween 80]/ [BSA NP] = 1.8 e7), is represented by a blue shift from 335 nm to 330 nm and a decrease in the fluorescence emission of 8 %. Again, these changes take place not showing changes neither in the hydrophobic character of the microenvironment surrounding the Trp_BSA NP_ nor in the exposure of the amino acid to the solvent (Figure 4.2f).

As the Tween 80 concentration increased no modifications in the structure of the BSA NP takes place. The blue shift that takes place throughout the experiment does not change the hydrophobic character of the TrpBSA NP binding pocket significantly. The same can be said for the amino acid exposure to the solvent (Figure 4.2e).

The BSA NP behaviour in the presence of chaotropic agents was compared to the BSA behaviour in the presence of chaotropic agents. Depending on the chaotropic agent, differences between the behaviours of the NP and BSA were observed. Tween 80 affected the BSA NP in a lesser way than it affected the BSA molecules (Figure 4.2). The effect this chaotropic agent has on the BSA NP can be explained using their size and charges. As Tween is a large molecule, it has difficulties accessing the NP, impeding any further denaturation of the inner core of the BSA NP. The charge of the molecule also contributes to the denaturation process. Tween 80 is a non-ionic detergent, summing this feature with its size, it is more difficult for the detergent to break through the NP and alter its structure after a superficial layer; the BSA NP may have less charged groups to its surface, making it harder for the detergent to interact with it, hence the less structure alteration. With the urea denaturation curve, it was observed that the Trp_BSA NP_ microenvironment barely increases its hydrophobicity, but the BSA NP structure experiences a rearrangement as the Trp is less exposed to the surface (35 % decrease of fluorescence emission) (Figure 4.2g). The decrease of the fluorescence emission can be related with the repulsion of the particle to the ionic groups of the surfactant in increasing concentration [10][34].

Contrary to this, the effect over BSA NP that the SDS has may be linked to the size, charge and penetration power. SDS is a relatively small molecule compared to Tween 80. Coincidentally, is the one who managed to change the BSA NP’s structure the most SDS manages to change the hydrophobicity in the Trp_BSA NP_’s vicinity as well as the amino acid exposure. Being an ionic chaotropic agent has a higher capacity of altering the structure of the NP. Thus, not only does its size help to get to the inner core of the BSA NP but also its charges enable it to destabilise the surroundings altering the NP whole structure.

According to the results, drugs dissolved in either urea or Tween 80 would be able to bind to the BSA NP without any significant alteration. In both media, the NP did not show structure denaturation degrees, which translate in the preservation of the nano vehicle’s functionality. Urea slightly affects the hydrophobicity of the Trp_BSA NP_ binding pocket, increasing this characteristic. It is known that hydrophobic drugs are prone to hydrophobic environments, which if this were true, then the increased hydrophobicity in the NP’s binding pocket may pose a higher affinity for the drug. As Tween 80 does not change the Trp_BSA NP_ surroundings, no modification in drug affinity should be noted. Nonetheless, SDS did affect the structure and surroundings of the Trp_BSA NP_, which results in a non suitable media for the NP as it may alter its functionality.

### 3.7 Ionic Strength

If IV administration is considered, another major feature to pay attention to is the stability of the BSA NP at different ionic strength levels. Even though both intracellular and extracellular environments have less than 2 M salt concentration [41], experiments were carried out using this concentration as the maximum. It is of great importance for the nanoparticle not to suffer any structure alteration once in these environments as its functionality might be affected, resulting in a drawback for its use as a drug delivery system. Therefore, the BSA NP was subjected to two different solutions: KCl and NaCl, in order to study its behaviour.

The ionic strength stability of the BSA NP in these solutions did not present significant protein precipitation (Figure 4.3a-b). We tried to cover a wide range of denaturing conditions to have a more detailed study; the BSA NP did not show significant protein precipitation (structure alteration) in neither of the conditions tested. Therefore, the BSA NP is stable in both intracellular and extracellular environment, so it should not present structure alteration. Hence, there would be no function lost before reaching the targeted site.

### 3.8 NP-Stability of time deviant behaviour

Knowing how stable the BSA NP is during a period is extremely important; alteration of the structure or agglomeration of the NP may affect its functionality. How the NP is affected by storage-time is, therefore, a significant concern. Samples in buffer 30 mM, PBS pH 7.0 were left under 4 °C and tested at different days (0, 15, 30, 60 days) by D.L.S, and fluorescence and FT – IR spectroscopy in search of possible structure alteration.

When measured at day 0, the BSA NP had a diameter size value of 71.53 nm. By the endpoint of the experiment, the diameter size of the BSA NP resulted in a 100 fold of its original size; 7359.7 nm (Figure 5a): the NPs aggregate among each other to form more prominent aggregates. The increase was first observed 15 days after the start of the experiment, where the diameter size was of 118.5 nm, doubling the original size. By day 30, the size obtained from measurements quadruplicate the original size c.a. 258.16 nm.

**Figure 5.**
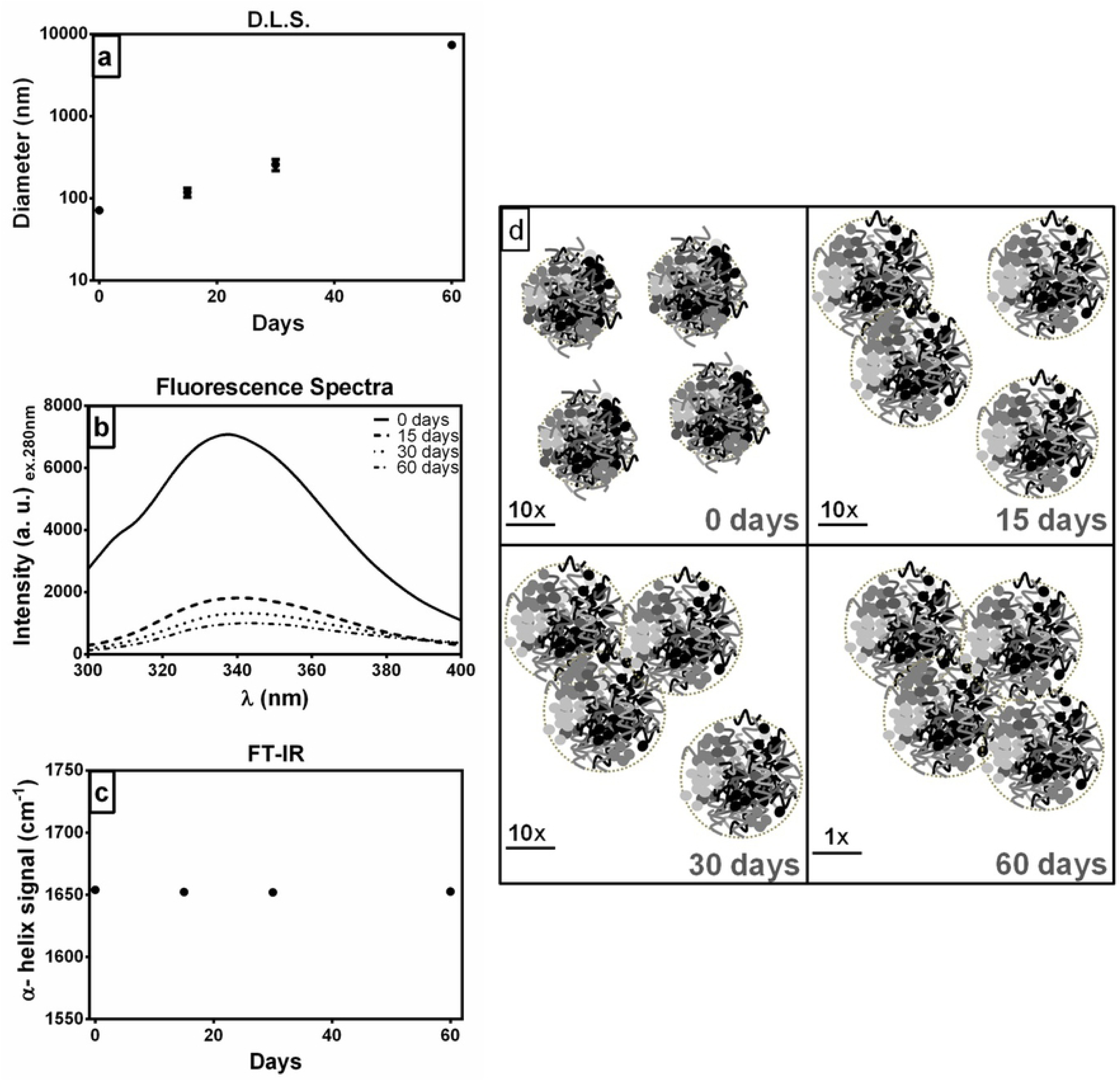
BSA NP stability in time measured by (a) Dynamic Light Scattering (DLS); (b) fluorescence spectra of the NP in the different days; (c) FTIR representing the α-helix signal; (d) Scheme representation of the BSA NP time behaviour stability.

By fluorescence spectroscopy, it was possible to observe not wavelength displacements, but changes in the fluorescence emission (Figure 5b). While comparing the fluorescence spectra of the BSA NP after the experiment started with the one obtained at the beginning of the experiment, we observed a decrease of fluorescence intensity emission of 75 % (day 15), 82 % (day 30), and 87 % (day 60). The decrease in the emission is taken as less Trp_BSA NP_ exposure to the solvent [10]. As there is no wavelength displacement, the microenvironment surrounding the Trp_BSA NP_ remains unchanged. Therefore, the decrease in the Trp fluorescence emission can be given by the aggregation of the NP; as they interact with one another, the Trp in each BSA NP is quenched by the NP coupled next to it and so on (Figure 5b).

When carrying – out the FT-IR experiment, the focus was on the α-helix maximum set at 1654 cm^−1^ (Figure 5c). This maximum describes the alteration, if any, in the global structure of the NP depending on its environment. A displacement to lower frequencies at day 15 to 1652 cm^−1^ takes place. This displacement stands for less α-helix group movements that emit the signal, seconding NP aggregation. Throughout the rest of the experiment, α – helix maximum remained at 1652 cm^−1^ (Figure 5c). This result backs up why no visible change in the BSA NP besides aggregation was observed in fluorescence spectroscopy (Figure 5b). Therefore, alteration of the structure of the BSA NP is observed at day 15, but no other structure alteration occurs after this point, only aggregation (Figure 5d).

As regards time stability, the BSA NP is preserved its native structure up to 15 days in 30 mM PBS pH 7.0, stored at 4 °C. More extended storage periods in these conditions may cause the colloidal suspension to flocculate and aggregate possibly altering its physicochemical characteristics. Therefore, an efficient way of long-term storage is highly recommended. In previous works, we have studied lyophilisation with trehalose as a way to preserve the structure/function of the NP without altering its structure or function [4].

### 3.9. Folic Acid attachment to the BSA NP

One of the advantages of the BSA NP is the number of functional groups. According to Lee et al., 2009, [41] no further modifications should be needed in order to enhance the NP functionality. For instance, sulphydryl groups are sometimes attached in order to target the BNP with thiol-selective reagents selectively [41]. The possibilities of ligands are such that they could decide the function of the NP for imaging, drug delivery or any other use, providing a more amicable NP to work with than the synthetic manufactured [41].

In our study, we propose the decoration of the BSA NP into folate coated BSA NP (FA-BSA NP). Folic acid was chosen due to its capacity to enhance the specificity of any NP. Moreover, its decoration to the NP is reproducible and low-cost. Cancer treatment has developed into two different ways; (i) the use of molecules to act as agents blocking protein expression pathways, or agents that are overexpressed in malignant cells, and (ii) the use of ligand which is compatible with an overexpressed receptor in a malignant cell [42].

We proposed two decoration models; decoration of the already irradiated BSA NP with folic acid, and decoration of BSA molecules with folic acid and then irradiate the samples for the production of the BSA NP.

The yield of each NP was estimated by either UV-visible (FA-BSA NP) (Figure 6.2a-c). The signal given by each NP was the information quantified for the yield of each process (Table 2). For FA-BSA NP the absorbance was measured at 352 nm (maximum for the FA in the BSA NP) (Figure 6.2b). The absorbance profile of this NP was different from the one of BSA NP as it showed a shoulder after 280 nm. The FA-BSA NP dispersion is free from non-attached FA polymers as it was separated by the size exclusion column before the study. Therefore, the shoulder in the absorbance profile refers to the attached FA to the BSA molecules forming the FA-BSA NP. Calculi carried out state that there are 3.07E+03 FA molecules absorbed for every BSA NP in the dispersion when the ligand is attached after irradiation of the BSA NP (Table 2). When the FA is attached to the BSA before the NP preparation process, the decoration renders 1.56E+02 FA molecules for every BSA NP (Table 2). The values refer to the FA absorbing at this wavelength; FA that is placed near or on the surface of the NP. Based on the calculi using the molecular weight of the BSA NP, there is more of the ligand present in the FA-BSA NPir than in the FA-BSA NP (Table 2). During the irradiation process, some FA polymers might be broken as well as some FA molecules might be hidden inside the formed NP resulting in fewer FA molecules per BSA NP detected.

**Figure 6.**
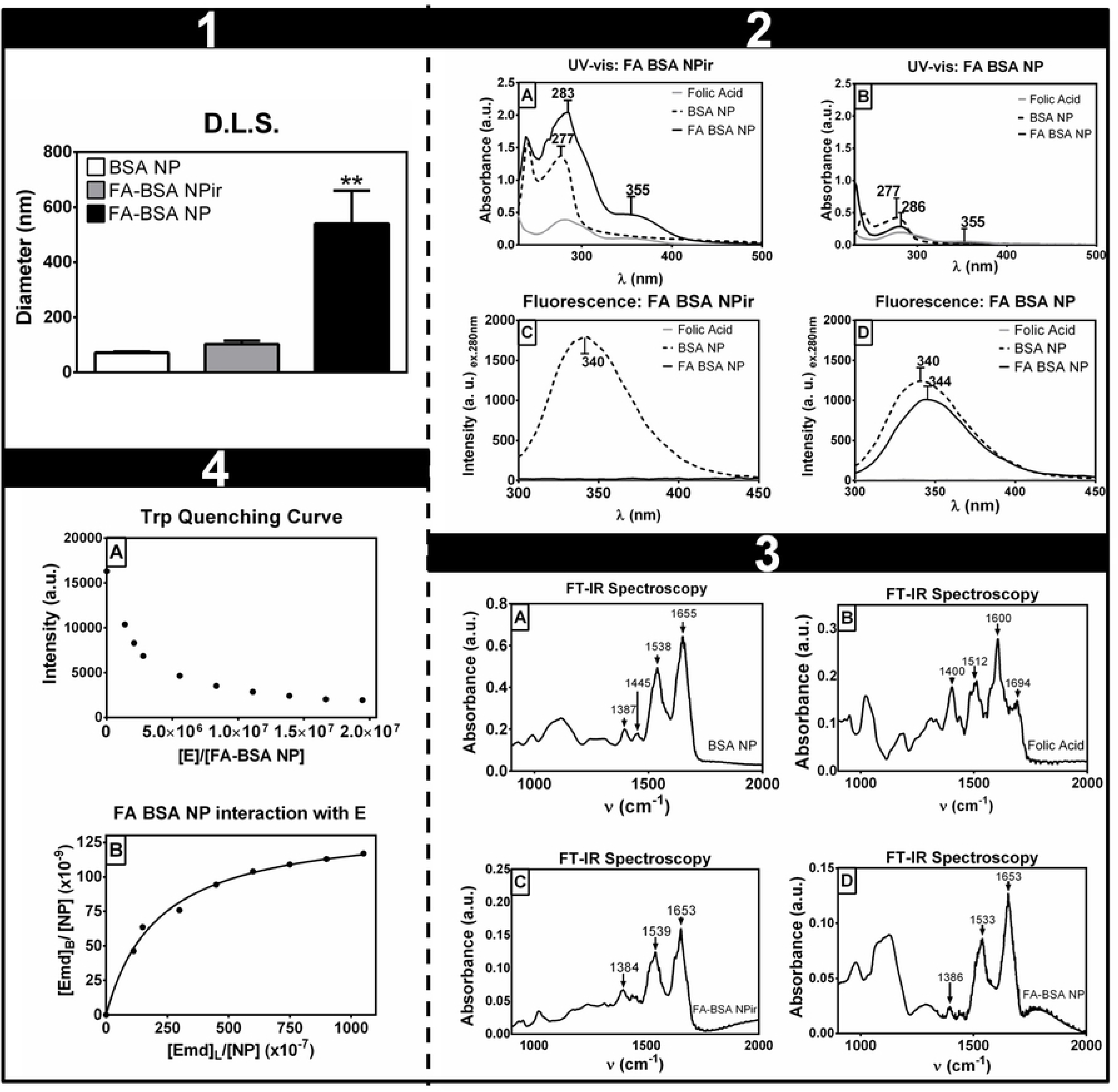
Characterisation of the NPs decorated with folic acid: 1 D.L.S study of BSA NP, FA-BSA NPir and FA-BSA NP in 30 mM PBS, pH 7.0 dispersions. 2 (A-B) UV-visible spectroscopy of the FA-NPs. Absorbance profile spectra go from 270-500 nm. (C-D) Fluorescence spectroscopy of BSA NP, FA-BSA NPir and FA-BSA NP where the fluorescence emission profile is between 300 – 450 nm, exciting the tryptophan in the NPs. 3 An FT-IR spectroscopy study of (A) BSA NP, (B) FA, (C) FA-BSA NPir and (D) FA-BSA NP; Absorbance profile go from 900 – 2000 cm^−1^. In every figure: BSA NP (dotted line), FA-NP (full line) and FA (gray line). 4 Functionality study of FA-BSA NP as a drug delivery system bound to hydrophobic drug E. (A) Quenching curve of the tryptophan fluorescence emission as E concentration increased. (B) Mathematical fitting was performed with the Scatchard equation [11].

**Table 2.** Approximation of the total ligand molecules per NP based on the experimental hydrophobic diameter.

| [BSA NP]<br>(mg/ml) | MW (Da) | NP<br>molecules/ml | [Ligand]<br>(mg/ml) | MW (Da) | Ligand<br>molecules/ml | Ligand/<br>BSA NP |
| --- | --- | --- | --- | --- | --- | --- |
| 0,34 | 4,71E+08 | 3,89E+17 | 2,75E-01 | 4,41E+02 | 3,75E+20 | 9,65E+05 |
| 0,043 | 1,58E+10 | 1,64E+15 | 1,40E-02 | 4,41E+02 | 1,91E+19 | 1,17E+07 |

#### 3.9.1. Characterisation of the FA-NPs:D.L.S

After each procedure, the hydrodynamic diameter was measured on each sample by dynamic light scattering (Figure 6.1). Results showed an increase in the diameter of the NP as AF was adsorbed to the surface; this confirms the existence of a new kind of BSA NP. The diameter size of the BSA NP decorated after gamma irradiation (FA-BSA NPir) was 102.90 ± 25.21, an increase of 144 % against the BSA NP size (71.53 ± 6.66 nm). The BSA NP decorated before gamma irradiation (FA-BSA NP) NP on its part, presents a hydrodynamic diameter of 332.1 ± 95.56 nm, the nanoparticle increases both its size (464 %), and its polidispersity as FA-BSA NPir.

Changes in the NPs size suggest changes in its structure. BSA NP size population is rather monodispersed, whereas the decorated samples seemed to be composed by more than one size population. The polydispersity increase in the FA-BSA NP is due to the aleatory and heterogenic character of the FA attachment process; the dispersion may contain naked BSA NP and BSA NP with FA attached to them (FA-BSA NP). Moreover, the FA tends to form polymers during its activation process with EDCI and NH, resulting in FA chains that will later be attached to the BSA [22]. These FA chains are the reason for the high polydispersity in the sample.

Based on this new piece of information, the molecular weight of both the FA-BSA NP was estimated, and correction on the yield of the preparation process was carried out (Table 3). We considered negligible the molecular weight of the FA molecule compared to that of the NP, and therefore, not taken it into account. The NPs could be embedded in a sphere where the volume of it would be given by, V=4/3 π R^3^. Once the volume was estimated, the mass of each NP was calculated by δ= m/ V, where the density (δ) value used was that of the BSA: 1.37 g/cm^3^. If we multiply the m value obtained by the Avogrado’s number, the molecular weight of each nanoparticle (MW) is given. This value when divided by the MW of the BSA molecule, gives the amount of BSA molecules per NP (Table 3).

**Table 3.**
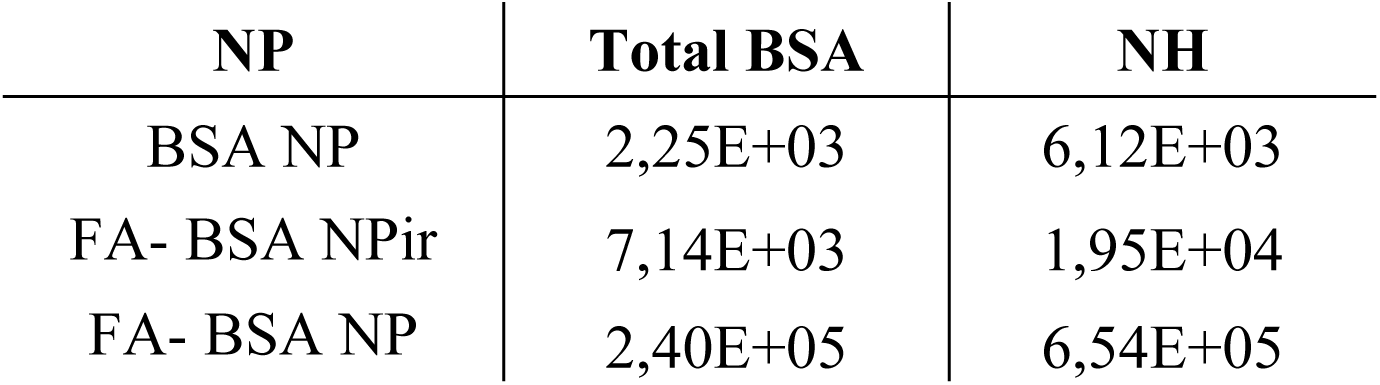
Total –NH surface groups per NP.

Based on these estimations, a reformulation on the yield of each NP preparation process was in order. According to the FA-BSA NPir MW (4.71 E08 g/mol), an ml of the dispersion contains 3.89 E17 NPs and 3.75 E20 FA molecules, giving an average of 9.65 E05 FA molecules/ NP (Table 3). As for the F-BSA NP MW (1.58 E10 g/mol), an ml of the dispersion contains 1.64 E15 NP molecules and 1.91 E19 FITC molecules. This results in 1.17 E73 FITC molecules/ NP (Table 3).

Theoretically, each FA-BSA NPir molecule is formed by 7.14 E03 albumin molecules, which would result in 1.95 E04 NH groups (for every BSA molecule there is 30 NH groups). As it is, each FA-BSA NP molecule is formed by 2.4 E05 albumin molecules, which would theoretically result in 6.54 E05 NH groups. If this were so, then the totality of the NH groups present in the NP would be attached to an FA molecule or FA polymer (Table 4).

**Table 4.** Interaction parameters of the FA-BSA NPE bioconjugate based on the values taken from the mathematical fitting in Figure 2b.

| Parameters | FA-BSA NPE |
| --- | --- |
| $B_{\text{mx}}$ | $1,4 \text{ e-}07 \pm 4,2 \text{ e-}09$ |
| $K_d$ | $2,1 \text{ e-}05 \pm 2,1 \text{ e-}06$ |
| $R^2$ | 0,99 |

#### 3.9.2. Characterisation of the FA-NPs: UV-visible

The absorbance profile of both NPs present wavelength displacement in their absorbance profile compared to that of the BSA NP (Figure 6.2a-b). The displacement represents a different hydrophobic character, which with the change in the width of the shoulder denote a change in the protein molecules structural change while in the NP. A change in the shoulder width refers to a change in the NPs size [23]; as the width increase, so does the size of the NP. The BSA NP has a shoulder width of 33 nm. The FA-BSA NPir has a shoulder width of 32 nm (1 nm thinner), which infers almost no change increase in size (Figure 6.2a). Contrary to this, the FA-BSA NP presents an increase in the shoulder width of 4 nm (c.a. 36 nm), which indicates a significant increase in the NPs size (Figures 6.2b). Results obtained in this experiment are consistent to those obtained in the D.L.S measurements (Figure 6.1).

#### 3.9.3. Characterisation of the FA-NPs: Fluorescence

The fluorescence emission of the tryptophan in each NP was obtained and the emission spectra compared (Figure 6.2c-d). The maximum for the BSA NP was set at 340 nm, while the FA-BSA NPir did not present any maximum in its emission spectra (Figure 6.2c). It is suggested that the reason for this is a quenching of the Trp because of the FA polymers proximity. For the quenching effect to take place, it is necessary a distance of 10 A; therefore, if this is true, then the FA will prevent the Trp fluorescence emission, blocking the hydrophobic pocket where the aromatic group is in. This possible situation would prevent any interaction of the NP with any given drug, which would result in a non-functional BSA NP. When the FA molecules are attached before NP preparation, the fluorescence spectra present a wavelength displacement to the red (maximum set at 344 nm), which indicates a decreased hydrophobic environment for the tryptophan in the NPs (Figure 6.2d) [10]. FA-BSA NP also presents a decreased intensity fluorescence emission, suggesting that the Trp in the FA-BSA NP is less exposed to the NP surface regarding the BSA NP (Figure 6.2d).

#### 3.9.4. Characterisation of the FA-NPs: FT-IR

For a better understanding of the interaction between the BSA NP and FA, an FT-IR spectroscopy experiment was carried out. For this particular analysis, we focused our attention on comparing the main signal maxima the BSA NP spectra showed and how this changed when the FA was present (Figure 6.3). In both samples, FA-BSA NPir (Figure 6.3c) and FA-BSA NP (Figure 6.3d), when the FA was present, the α-helix signal experienced a frequency displacement from 1655 – 1653 cm^−1^; this displacement to lower frequency waves mean less freedom of movement, assuming that a more compact structure as this maximum represents the global structure of the BSA NP. The β-sheet signal, another significant maximum for the NP structure, shows a displacement to higher frequency waves in the FA-BSA NPir and lower ones for FA-BSA NP; from 1538 cm^−1^ (BSA NP) to 1539 cm^−1^ (FA-BSA NPir) and 1533 cm^−1^ (FA-BSA NP). While in the BSA NP decorated after irradiation the FA did not do a significant alteration in its β-sheet composition, in the BSA NP decorated before irradiation the FA molecules seem to have organised the β-sheet present in the NP in a more compact way. The interaction with the FA molecules is given by –NH present in the structure of the albumin molecules forming the different NPs. When added to the sample before irradiation, the FA molecules arrange themselves in a way that makes the β-sheet structure more compact, which may also mean that these molecules are also involved in the spatial arrangement of BSA molecules forming the NP. On the contrary, when added after the NP is formed, the FA molecules are only interacting with the –NH exposed to the surface, altering in a minor way the β-sheet structure arrangement but not the global structure of the NP (Figure 6.3).

The biophysical characterisation of the FA-NPs showed that the most efficient method for FA decoration of the BSA NP is to attach the ligand to the BSA molecules before NP preparation. Otherwise, the polymers of the FA are at risk of interacting with the Trp of the BSA NP preventing any drug to bind in the main NP binding pocket (Figure 6.2c). As it is, functionality studies were carried out with the FA-BSA NP, as the NP showed no interfering of the FA molecules with the main hydrophobic pocket of the NP.

#### 3.9.5. Interaction study: FA-BSA NP with Emodin (E)

In order to study the FA-BSA NP functionality as a drug delivery system, its interaction with the hydrophobic drug emodin (E) was studied (Figure 6.4). E is has shown in recent years a potential antitumor effect and attempts have been made to diminish its secondary effects by using different drug delivery systems. It was chosen for this study as it also presents the ability to bind to proteins forming complexes [11].

A fluorescence study was carried out in order to estimate the interaction parameters such as dissociation constant (Kd) and maxima drug loading capacity (Bmx) [11] (Figure 6.4). The interaction between the FA-BSA NP and E was described by the quenching curve of the tryptophan fluorescence emission of the NP when being incubated with increasing concentrations of E (Figure 6.4a). From the data collected in the experiment, an interaction curve can be represented (Figure 6.4b), where a mathematic fitting is done in order to obtain the interaction parameters (Table 5). If Kd is a significant value, then the association constant (Ka) should be small. Therefore, larger values of Kd suggest a low affinity between the NP and drug [11][43]. As it is, FA-BSA NP presents a low value of Kd, which means there is an excellent affinity for E (Table 5).

According to Sevilla et al., 2007 [11], the Kd for BSA and E varies between 10 e-08 and 10 e-06. In this work the Kd obtained is in the order 10 e-05, which suggest the affinity is increased because of the derivatisation of the NP. During the preparation process (FA attachment to the BSA and following irradiation), the FA molecules and polymers generate a dense aggregation of the NP. This solid aggregation results in protein structure alteration, increasing the affinity for the E despite the decreased hydrophobic character of the binding pocket (Figure 6.4).

#### 3.9.6. Cytotoxicity study in MCF-7 cells treated with FA-BSA NP

The cytotoxic effect of the FA-BSA NP and the bioconjugate FA-BSA NPE was studied by the effect they had on the cell metabolic activity in the human tumour breast cell line MCF-7. The effects were studied up to 48 hs post incubation, maximum expression of the E effect [44] (Figure 7).

**Figure 7.**
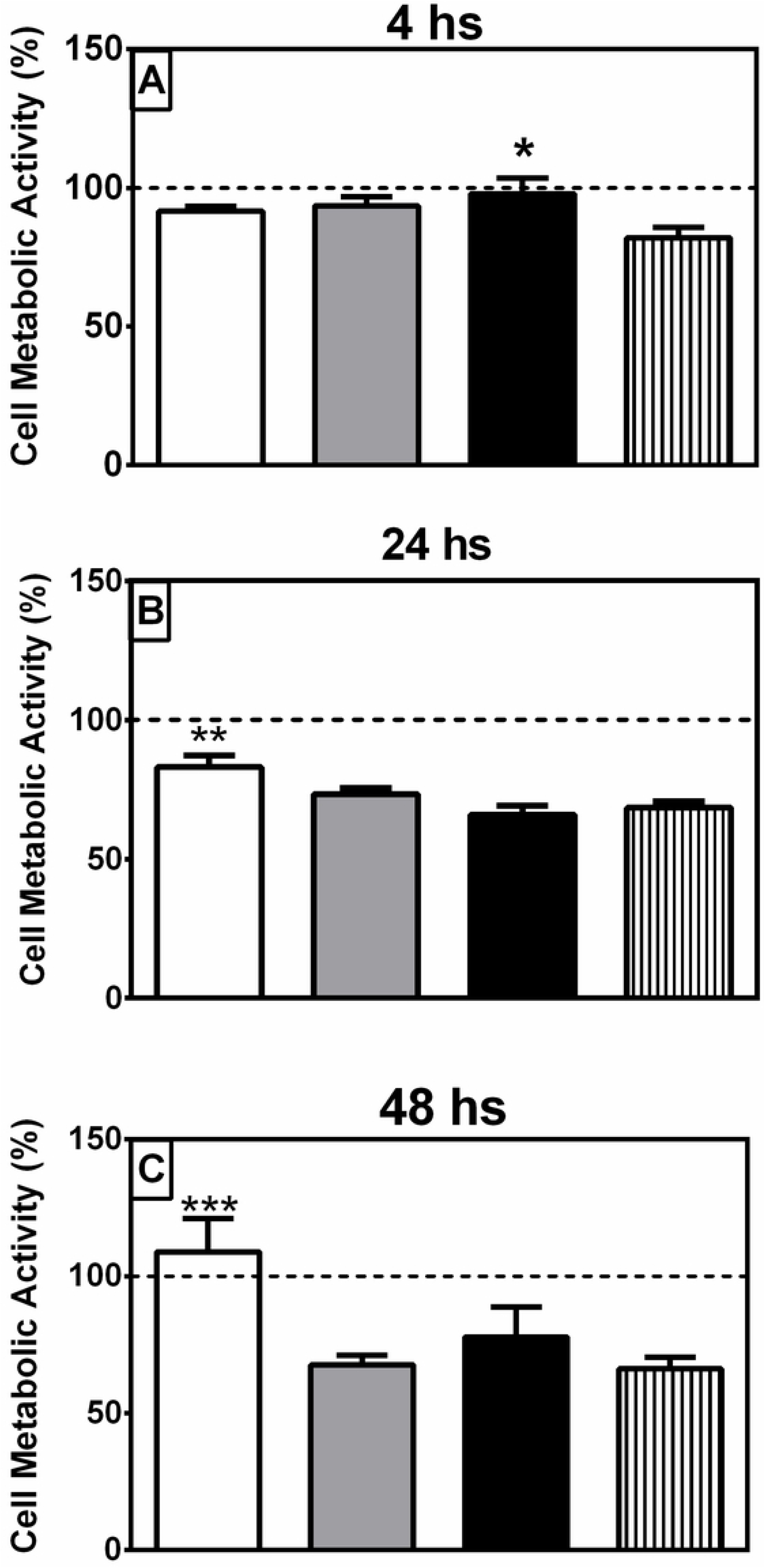
Cytotoxic study in MCF-7 cell line by an MTT test at 4, 24 and 48 h post-FA-BSA NP, FA-BSA NPE, and free E incubation. The 100 % dot line refers to cells without treatment.

After 4 h of incubation with the different treatments, cells treated with the bioconjugate and FA-BSA NP generate a 10 % decrease in the cell metabolic activity, whereas the cells treated with E generate a 15 % decrease in the cell metabolic activity (Figure 7). At 24 h post incubation, the metabolic activities for cells treated with the bioconjugate and with free E decrease 30 %, while the metabolic activity for cells treated with FA-BSA NP presents a lesser decrease (15 %). At 48 h post incubation, the cell metabolic activity reaches only 55 % when treated with the bioconjugate and with free E. The FA-BSA NP does not alter the cell metabolic activity 48 h post incubation (Figure 7).

The cytotoxicity study suggests that while FA-BSA NP is not toxic for the treated cells, the bioconjugate can decrease the cell metabolic activity similarly than the free E can do. Results encourage thinking of the FA-BSA NP as a nanoparticle with great potential as a drug delivery system.

### 3.10. Cell immune secretion by NPs

Every cell treated with the different stimulants presented secretion of the transcription factor NFκB (Figure 8). No differences were found among the stimulants BSA, BSA NP y FA-BSA NP (Figure 8a). Every one of them presents an inflammatory dose-dependent response between 80 – 5.5 nM. At the highest concentration of each sample, the immune response doubles that of the basal response (cells without treatment).

**Figure 8.**
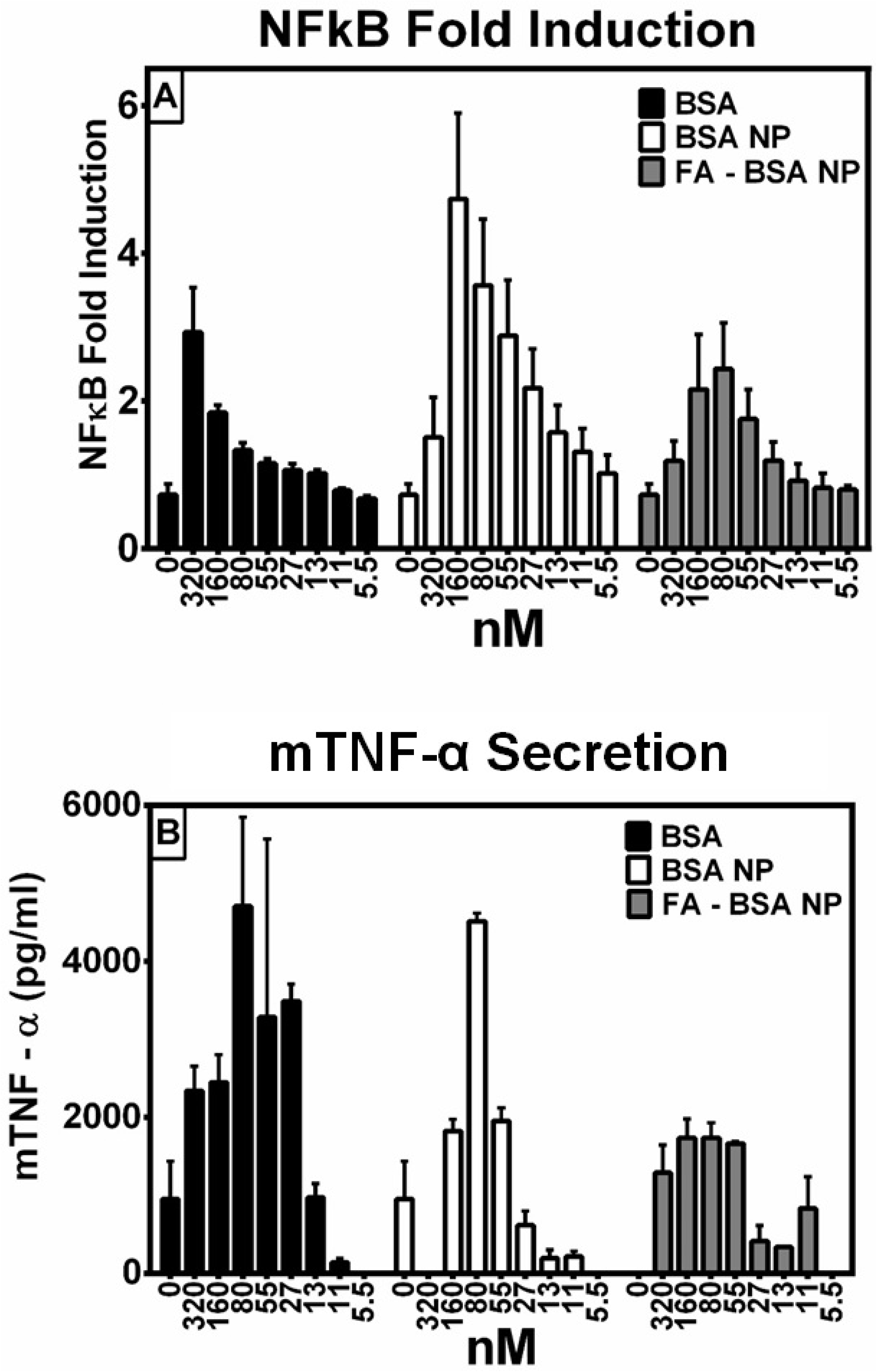
Cell immune response assay based on the BSA NP and FA-BSA NP stimuli. (A) NFkB expression of the treated cells. (B) TNF-Secretion of the cells treated. Cells were also stimuli with molecular BSA for further comparison

The secretion TNF – α was also tested, showing secretion in every cell tested, independently from the stimulant incubated (Figure 8b). As the both TNF – α and NFκB are involved in the same signaling pathway; it is not surprising that results are in agreement with each other.

It is believed that there is a connection between the β – sheet percentage in the NP with the secretion of a pro-inflammatory response in the cell. The β – sheet structures are involved in the forming of amyloid-like fibers, toxic to the immune system. From the FT-IR study carried out in the different NPs here tested, we were able to observe the presence of such structures in every one of the NPs. These described structures may be the cause of the immune cell response. NPs, in general, are capable of generating two different types of immune response: (1) they can stimulate it, acting as adjuvants, (2) they can be toxic for the organism [45]. Both, size and physic-chemical structure are responsible for this.

Before this study, we expected the BSA NP no to generate a toxic effect on the cells tested as BSA molecules form it and albumin is not a strange molecule for the organism. Nevertheless, Wheeler et al., 2011 [46], stated that the albumin is capable of pro-inflammatory effects in the cell, described by an expression of the transcription factor NFκB. In our study, molecular albumin has an immune response in a dose-dependent manner (Figure 8). This trend and values are similar when the cells were stimulated with BSA NP. Both samples double the immune response compared to the basal response in their highest concentration, despite the differences in the molecule diameters (BSA 6 nm and BSA NP 70 nm). The FA-BSA NP stimulated cells do not show a high immune response. This NP also generates a dose-dependent immune response, even though the response obtained is lower than the other NP.

Contrary to Wheeler et al., 2011 [46], in this study a higher diameter (FA-BSA NP 332 nm), does not mean a higher immune response. The reason behind this may be the presence of FA in the NP. This molecule acts as a ligand of macrophages, which might mean the immune cells recognize the FA-BSA NP without the necessity of generating a pro-inflammatory response.

## 4.0 Conclusion

It was possible to characterise a crosslinked γ-irradiated albumin nanoparticle as a potential novel nanovehicle for drug delivery. Preservation of the primary function of the albumin was observed with enhanced binding efficiency once the NP was formed, as well as an enhanced drug release kinetic profile. The improvements shown by the NP have wanted features in the upcoming field of medicine, thus encouraging the use of the BSA NP in ongoing research.

The BSA NP was tested against different adverse experimental conditions for its stability. When tested in extreme acidic solutions (pH 2.0), the BSA NP adopts a denatured state similar to molecular BSA. In this condition, the NP has an extended structure where the longitudinal dimensions are maximised. At pH 9.0, a certain level of NP denaturation was observed, even though this conformation state was not comparable as the one at pH 2.0. BSA NP behaviour with the different chaotropic agents resulted in discriminating the drugs to be bound to the NP. Those dissolved in urea and Tween 80 would be a suitable choice, but no those drug in need to be dissolved in SDS, as this media denatures the NP structure. A highlight in this section of the work was to decipher some characteristics of the NP structure: it stability in urea and Tween 80 is a consequence of a more compact structure that gives protection against said chaotropic agents. As far as ionic strength stability goes, the nanoparticle proved to be in its native form up to 2 M solutions, predicting high stability in both intra – and extracellular environments.

The highlight of the whole study relays in the drug release profile of the BSA NP at pH values similar to the tumour ones. The BSA NP at pH 5.5 – 6.0, released around 40 % of E after 24 hours. The BSA NP had no toxic effect on cancerous or blood cells by itself, proving to be innocuous. On the other hand, when bound to E, nonetheless the bioconjugate reached 50 % of cell mortality in cancerous cells such as MCF7 and PC-3.

The BSA NP is a suitable nano-delivery system for hydrophobic antitumoral drugs, maintaining the original molecular properties of the protein but enhancing its delivery features. It has so far proven to be successful against cancerous cells, but innocuous to healthy human blood cells. Results in this work highlight the potentiality of the NP as a nanovehicle, encouraging future research on it.

## 5.0 Acknowledgements

This work was supported by grants from Consejo Nacional de Investigaciones Científicas y Técnicas, Universidad Nacional de Quilmes, Ministerio de Ciencia y Tecnología (MINCyT), Comisión de Investigaciones Científicas de la Provincia de Buenos Aires, Nuclear Atomic Energy Agency (IAEA) CRP codes F220064 and F23028.

We are also thankful to Dr. Beatriz Patricio from the Institute of Biophysics, Universidade Federal do Rio de Janeiro, Rio de Janeiro, Brazil and to Dr. Constanza Flores from Laboratorio de Materiales Biotecnológicos from Universidad Nacional de Quilmes, Bernal, Quilmes, Buenos Aires, Argentina for help with the AFM microscopies.

## Author Contributions

The manuscript was written through the contributions of all authors. All authors have approved the final version of the manuscript.

